# Cholinergic modulation engages a CHRM1-IL-17 neuroimmune axis to promote diabetic wound repair

**DOI:** 10.64898/2026.11.06.764960

**Authors:** Zhuqing Li, Georgios Theocharidis, Sangmin Lee, Nikolaos Kalavros, Sheethal Umesh Nagalakshmi, Mauricio Contreras, Eleftheria-Angeliki Valsami, Hyangsu Nam, Ming Guan, Guangyu Chu, Ikram Mezghani, Enya Wang, Brandon J Sumpio, Jinfei Hou, Matheus Gennari-Felipe, Jessica Gilman, Nuria Lafuente-Gómez, Lidia Maeso Aragón, Nikolaos Dimitrakakis, Antonella Arruda de Amaral, Shuoshuo Wang, Ioannis S Vlachos, David J Mooney, Aristidis Veves

**Author notes:** These authors contributed equally (Zhuqing Li, Georgios Theocharidis, Sangmin Lee).

## Abstract

Chronic non-healing wounds are a major complication of diabetes, where persistent inflammation fails to transition into effective tissue repair (*1–3*). Here we identify cholinergic modulation as a mechanism that promotes productive inflammation and diabetic wound healing. Using a healing-associated fibroblast signature derived from single-cell profiling of human diabetic foot ulcers (*3*), we identified the acetylcholinesterase (AChE) inhibitor metrifonate as a candidate inducer of reparative wound programs. Topical delivery accelerated healing in diabetic mice and increased acetylcholine availability within the wound. Complementary receptor pharmacology identified muscarinic acetylcholine receptor 1 (CHRM1) as the principal receptor mediating the reparative response. Transcriptomic and cellular analyses revealed early induction of IL-17-associated inflammatory programs and enrichment of CHRM1⁺NK1.1⁺ innate lymphocytes. Depletion of NK1.1⁺ cells or neutralization of IL-17A abolished the pro-healing effect, whereas spatial transcriptomics revealed treatment-associated organization of NK cells within vascular niches. Localized treatment further promoted repair in diabetic porcine wounds and human skin models. Together, these findings identify a CHRM1-IL-17 neuroimmune axis linking cholinergic modulation to diabetic wound repair and establish local AChE inhibition as a potential therapeutic strategy for chronic wounds.

---

Diabetic foot ulcers (DFUs) are among the most devastating complications of diabetes and remain a leading cause of lower-limb amputation worldwide (*1, 4*). Despite advances in wound care, debridement strategies, infection control, and pressure offloading, the majority of therapies target the wound itself rather than the biological mechanisms that govern successful tissue repair (*5, C*). As a result, healing remains slow, incomplete, and highly variable among patients, underscoring the need to identify fundamental pathways capable of restoring effective repair in the diabetic wound microenvironment (*7*).

Successful wound healing requires tightly coordinated interactions between immune, stromal, vascular, and epithelial cells (*2, 8*). Increasing evidence suggests that chronic diabetic wounds reflect not only cell-intrinsic dysfunction, but also disruption of the multicellular programs that normally coordinate inflammation, tissue remodeling and regeneration (*S-12*). Single-cell and spatial transcriptomic technologies have transformed our understanding of this cellular complexity, revealing previously unrecognized cell states and intercellular signaling networks that distinguish healing from non-healing wounds (*3, 13–15*). However, how these systems-level insights can be translated into therapeutic strategies remains largely underexplored (*1C*).

Here, we leverage a healing-associated fibroblast transcriptional signature identified in our previously published single-cell atlas of human DFUs (*3*) to perform computational drug repurposing using Connectivity Map (CMap) (*17, 18*). This analysis nominated the acetylcholinesterase (AChE) inhibitor metrifonate as a leading candidate predicted to induce a pro-reparative transcriptional program. By integrating computational drug discovery with biomaterial engineering, multimodal molecular profiling, spatial transcriptomics, and functional immune perturbation across murine, porcine, and human wound models, we identify a previously unrecognized CHRM1-IL-17 neuroimmune axis that promotes diabetic wound repair.

## Metrifonate discovery and delivery

To identify therapeutic strategies capable of restoring impaired healing in DFUs, we leveraged our previously generated single-cell and spatial transcriptomic atlas of human DFUs, which identified a healing-associated fibroblast (HE-fibro) state selectively enriched in wounds that successfully healed. We hypothesized that pharmacological induction of this reparative transcriptional program could restore healing in chronic diabetic wounds.

We therefore compared HE-fibro with canonical fibroblasts and used the resulting differential-expression profile to identify compounds predicted to induce the HE-fibro state. ASGARD (*1S*) analysis of Connectivity Map/LINCS perturbational profiles identified metrifonate as the second-ranked compound among 11,908 perturbagens (**Fig. 1A,B** and **data S1**). Metrifonate is a clinically evaluated AChE inhibitor originally developed as an anthelmintic agent and previously proposed for the treatment of Alzheimer’s disease (*20, 21*).

To enable controlled topical delivery, we developed a metrifonate-loaded alginate hydrogel incorporating laponite clay nanoparticles to mediate ionic interactions with both metrifonate and the alginate matrix. Aqueous solutions of alginate, metrifonate, and laponite were cast into molds, lyophilized, and crosslinked to generate disc-shaped hydrogels (**Fig. 1C**). The resulting metrifonate-loaded hydrogels (Met-gels) exhibited a macroporous architecture with an average pore size of 14.4 ± 5 μm, high surface area and water retention, with minimal mass loss during incubation in PBS (**Fig. 1D** and **fig. S1A,B**). Incorporation of laponite was confirmed by Fourier transform infrared spectroscopy (**fig. S1C**). We next determined whether laponite could modulate metrifonate release. Alginate/laponite hydrogels sustained metrifonate release over five days, whereas hydrogels lacking laponite exhibited rapid drug release (**fig. S1D**). Increasing the drug dose at a fixed laponite concentration accelerated release, consistent with a limited capacity of the clay to retain metrifonate (**Fig. S1E, H, I**). Increasing laponite content in parallel with higher metrifonate doses restored sustained release and reduced the initial burst (**fig. S1F, G**). Based on these findings, three formulations exhibiting controlled release over five days were selected for subsequent studies: low-dose (8 μg metrifonate, 10 mg ml⁻¹ laponite), medium-dose (50 μg metrifonate, 30 mg ml⁻¹ laponite) and high-dose (200 μg metrifonate, 30 mg ml⁻¹ laponite) gels (**Fig. 1F**). The low-dose formulation released approximately 1 μg metrifonate per day for four days, yielding a near linear release profile over this interval (**Fig. 1F and fig. S1J**).

Finally, we assessed whether metrifonate retained its bioactivity following hydrogel storage. Met-gels stored at room temperature or -20 °C for 17 days released metrifonate over three days with AChE inhibitory activity comparable to that released from freshly prepared gels, demonstrating preservation of drug bioactivity under both storage conditions (**Fig. 1G**).

## Transient metrifonate treatment accelerates diabetic wound healing

To define the therapeutic window for localized metrifonate delivery, we evaluated multiple dosing regimens in the *db/db* mouse model of impaired diabetic wound healing (*22*). Based on the *in vitro* release profiles, which showed comparable drug release from the medium- and high-dose formulations by day 3, we focused on low- and medium-dose hydrogels with defined removal time points to determine the minimum duration of exposure required for therapeutic efficacy (**Fig. 2A**).

A low-dose Met-gel removed on day 3 consistently produced the greatest acceleration in wound closure, outperforming prolonged treatment and higher-dose regimens (**Fig. 2B-D and fig. S2A, B**). These findings demonstrate that metrifonate acts within a narrow early therapeutic window and that sustained drug exposure does not further enhance repair. Accordingly, the low-dose gel removed on day 3 was selected for all subsequent in vivo studies, with matched blank hydrogel controls.

To capture early mechanistic events preceding macroscopic wound closure, we introduced an earlier harvest time point at day 5 (**Fig. 2E**). At this stage, metrifonate treated wounds exhibited a trend for enhanced re-epithelialization (**Fig. 2G, J**), increased hyperproliferative epidermal area (**Fig. 2H**), and elevated angiogenesis compared to controls (**Fig. 2I, J**). RT-qPCR gene expression analysis revealed reduced expression of markers *Tgfb1* and *Acta2*, associated with fibroblast activation and tissue remodeling together with reduced expression of inflammatory mediator *Il1b* (**Fig. 2K and fig. S2I**) in treated wounds. By day 10, treatment-associated differences were largely attenuated across histological, vascular, and inflammatory/remodeling gene expression endpoints (**fig. S2C-F, K**). Tissue immunofluorescence analysis further demonstrated decreased neutrophil (NIMP-R14+ cells) numbers and increased macrophage (F4/80+ cells) accumulation in metrifonate treated wounds, consistent with accelerated progression of the inflammatory response (**Fig. 2N-P**). A down-regulation in protein levels of MMP-9, TIMP-1 and NGAL corroborates this finding (**Fig. 2L, M**) (*23*). Mast cell abundance and degranulation were likewise increased, in line with their previously described pro-reparative roles in diabetic wound healing (**fig. S2G, H)** (*24*).

Together, these findings demonstrate that a brief period of localized cholinergic modulation during the early inflammatory phase is sufficient to accelerate diabetic wound repair by rapidly reshaping the wound immune microenvironment.

## CHRM1 mediates cholinergic diabetic wound repair

To determine whether metrifonate-mediated wound repair results from sustained cholinergic signaling, we first quantified AChE activity and acetylcholine levels within the wound microenvironment. Despite Met-gel removal on day 3, AChE activity remained suppressed and acetylcholine levels remained elevated at day 5, confirming persistent local cholinergic modulation following transient treatment (**fig. S2K, L**).

To identify receptor candidates during the early stages of repair, wounds were harvested on days 1 and 2, dissociated into single-cell suspensions, and separated by magnetic-activated cell sorting (MACS) into neutrophil-enriched and non-neutrophil fractions prior to RNA isolation (**Fig. 3A**). This strategy was adopted because neutrophils constitute the predominant immune population during the initial inflammatory phase of wound healing (*25*), allowing cholinergic receptor expression to be assessed independently in early infiltrating neutrophils and the remaining wound cell populations. This separation strategy was applied consistently across subsequent receptor expression analyses and downstream transcriptomic studies. AChE inhibition induced dynamic, cell type-specific changes in cholinergic receptor expression during the first two days after injury. Multiple muscarinic and nicotinic receptors were transiently upregulated in both neutrophil-enriched (**Fig. 3B, D**) and non-neutrophil fractions (**Fig. 3C, E**), with *Chrm1*, *Chrm3*, and *Chrna7* showing the most consistent induction. These findings suggest that localized cholinergic modulation rapidly engages multiple receptor pathways during the early inflammatory response.

To determine which receptor was functionally required for therapeutic efficacy we next examined the effects of selective muscarinic and nicotinic receptor activation and blockade (**Fig. 3F-J**). Topical inhibition of CHRM1 completely abolished the pro-healing effects of metrifonate (**Fig. 3G**), whereas blockade of CHRM3 or α7 nicotinic receptors had no effect on wound closure (**Fig. 3H, I**). Conversely, selective activation of CHRM1 was sufficient to phenocopy metrifonate treatment, while CHRM3 and α7 nicotinic receptor agonists failed to improve healing (**Fig. 3J**).

To determine whether this therapeutic response was unique to metrifonate or reflected a broader consequence of AChE inhibition, we evaluated three additional clinically used AChE inhibitors. Rivastigmine, donepezil and galantamine each accelerated diabetic wound healing in *db/db* mice, demonstrating that the reparative response extends across structurally distinct AChE inhibitors rather than being unique to metrifonate (**fig. S3**).

Together, these findings identify CHRM1 as the principal receptor mediating the pro-reparative effects of localized cholinergic modulation and establish AChE inhibition as the upstream trigger of this response.

## Cholinergic modulation induces IL-17 in diabetic wounds

Having established CHRM1 as the principal receptor mediating metrifonate activity, we next characterized the downstream molecular programs engaged by cholinergic modulation during the early stages of repair. The MACS-separated neutrophil-enriched and non-neutrophil fractions generated for cholinergic receptor profiling (**Fig. 3**) were subjected to bulk RNA sequencing to define treatment-induced transcriptional responses during the first two days after injury (**Fig. 4A-E**).

Bulk RNA sequencing revealed a temporally coordinated and compartment-specific transcriptional response to cholinergic modulation during the early phase of wound healing. At day 1, transcriptional changes were limited, with the neutrophil-rich fraction showing induction of stress-and inflammatory-response genes, including *Hmox1* and *Cxcl10*, whereas the remaining wound cell fraction exhibited early induction of *Il1a*, *Inhba*, *Ccl3* and *Il1rn* (**fig. S4F** and **data S2**) (*2C, 27*). By day 2, this initially restricted response had expanded markedly in both fractions (**Fig. 4A, C** and **fig. S4A-E**). In the neutrophil-rich fraction, AChE inhibition induced a broad inflammatory program with 2495 upregulated transcripts, encompassing chemotactic and granulocyte-associated genes, including *Cxcl3*, *Csf3*, *Fpr1*, *Il1b* and *Nlrp3* (**Fig. 4A**) (*28, 2S*), and with enrichment of pathways related to inflammatory responses, granulocyte trafficking and IL-17 signaling (**Fig. 4B**) (*S, 30*).

In parallel, the non-neutrophil fraction also exhibited a robust upregulation of 499 genes and developed a prominent IL-17-associated lymphoid program, with *Il17a* emerging as the most strongly induced transcript and accompanied by *Il17f*, *Il7r*, and additional lymphoid-associated transcripts, together with *Il23a* and angiogenesis and tissue-remodeling programs (**Fig. 4C, D** and **data S2**) (*31, 32*). Temporal analysis further identified genes exhibiting opposing day 1-to-day 2 trajectories between control and treated wounds, consistent with treatment-dependent remodeling of the early inflammatory response (**Fig. 4E** and **data S3**). Together, these findings indicate that AChE inhibition does not simply increase inflammatory signaling at a single time point, but reshapes its early kinetics, progressing from a restricted day 1 response to a coordinated day 2 program characterized by IL-17-associated signaling and granulocyte-directed inflammatory effector pathways.

In accordance with this transcriptional response, multiplex analysis of day 3 wound lysates showed increased IL-17 protein together with IL-5, IL-7, IL-9 and a trend toward increased VEGF (**Fig. 4F-M** and **table S1**) (*33*). Thus, transcriptional emergence of an IL-17-associated inflammatory program by day 2 preceded and was reflected by increased cytokine abundance at day 3.

Collectively, these findings demonstrate that cholinergic modulation rapidly remodels the early wound inflammatory environment and identifies IL-17 signaling as a prominent downstream pathway associated with enhanced diabetic wound repair.

## NK1.1⁺ innate lymphocytes mediate diabetic wound repair

To identify the cellular drivers of metrifonate-mediated tissue repair, we first performed broad flow cytometric profiling across major myeloid and lymphoid lineages. Gating strategies were outlined in **fig. S5**. While overall leukocyte composition remained largely unchanged, apart from a modest change in abundance for monocyte-derived dendritic cells (DCs), natural killer (NK) cells (CD45⁺CD3ε⁻NK1.1⁺) emerged as the only population examined exhibiting significant increases in both the CHRM1⁺ and IL-17A⁺ fractions (**fig. S6A-C**). Immunofluorescence staining independently confirmed increased CHRM1⁺NCR1⁺ NK cells in day 3 wounds, whereas corresponding increases in CHRM1 expression were not detected in macrophages, DCs or mast cells (**fig. S6D-G**). Prompted by this unbiased screen, we focused on the NK1.1+ lymphoid compartment in the wound microenvironment. Flow cytometry and tissue immunofluorescence confirmed that AChE inhibition markedly increased CHRM1+and IL-17A+expressing NK1.1+ cells within the healing wound bed (**Fig. 5A-F**). Given that IL-17A expression in innate lymphoid populations is canonically regulated by the transcription factor RORγt (*34, 35*), we next investigated whether cholinergic stimulation drives a broader RORγt-mediated reparative program. In line with our IL-17A readouts, both flow cytometry and tissue immunostaining revealed a striking enrichment of double-positive RORγt+CHRM1+ and RORγt+IL-17A+ cells in LD Met-gel treated wounds (**fig. S7**). Because RORγt is expressed across multiple innate and innate-like lymphocyte populations (*3C*), and NK1.1 is shared by conventional NK cells and several innate-like T-cell subsets (*37*), we performed deeper phenotypic profiling to characterize the RORγt⁺ lymphocyte populations associated with the response. This analysis identified treatment-associated increases in RORγt⁺ mucosal-associated invariant T (MAIT) cells and invariant natural killer T (iNKT) cells, with both populations exhibiting increased expression of CHRM1 and IL-17A (**fig. S8A-M**). Despite these phenotypic responses, metrifonate retained its pro-reparative activity in both *Mr*1-/-(*38*) and *Traj*18-/-(*3S*) STZ-induced diabetic mice, indicating that neither MAIT nor iNKT cells are individually required for wound repair (**fig. S8N-O**). To determine whether these cellular and cytokine responses were functionally required for therapeutic efficacy, we performed targeted immune cell depletion and cytokine neutralization in the *db/db* model (**Fig. 5G**). Neutralization of IL-17A completely abolished the wound healing benefit of metrifonate, as did depletion of NK1.1⁺ cells. In contrast, depletion of γδ T cells or neutrophils did not impair the response to treatment, indicating that these populations were dispensable for its therapeutic efficacy (**Fig. 5H, I**).

Together, these findings implicate CHRM1-expressing NK1.1⁺ innate lymphocytes as critical cellular mediators of an IL-17A-dependent reparative response following localized cholinergic modulation.

## Cholinergic modulation shapes vascular-NK niches

To define how localized cholinergic modulation reorganizes the wound microenvironment, we performed Xenium high resolution spatial transcriptomic profiling of day 3 wounds using a custom designed 471-gene panel encompassing immune, stromal, vascular, epithelial, and neuroimmune signaling genes (**table S2**). Unsupervised clustering for a total of 237,213 cells resolved the major cellular populations within the wound and peri-wound areas, including neutrophils, macrophages, DCs, NK and T/NKT-like cells, fibroblast subsets, vascular endothelial cells, smooth muscle cells (SMCs), keratinocytes, and melanocytes, enabling spatial analysis of immune and stromal organization during repair (**Fig. 6A, B** and **fig. SG**). Cell type-specific pathway analysis demonstrated distinct spatial compartmentalization of cholinergic and downstream reparative signaling. CHRM1-associated signaling was enriched primarily within neutrophils and NK cells, consistent with preferential cholinergic responsiveness within these populations. In contrast, IL-17-associated transcriptional programs extended across multiple immune and stromal populations, with the strongest enrichment observed in neutrophils, NK cells, T/NKT-like cells, and SMCs (**Fig. 6C**), suggesting propagation of downstream reparative signaling throughout the wound microenvironment. Because NK1.1⁺ innate lymphocytes emerged as the critical cellular mediators of AChE inhibition-induced repair, we next investigated their spatial organization. Neighborhood enrichment analysis revealed preferential localization of NK cells adjacent to SMCs and perivascular fibroblasts within both the drug treated tissues’ wound edge and wound center (**Fig. 6D**), identifying previously unrecognized vascular-associated neuroimmune niches. Ligand-receptor inference analysis predicted enhanced bidirectional communication between NK cells and both SMCs and vascular endothelial cells, particularly at the wound edge (**Fig. 6E**). These analyses nominate the wound edge vasculature as a potential site of enhanced NK-vascular signaling following cholinergic modulation.

Together, these findings reveal that localized cholinergic modulation reorganizes the diabetic wound into spatially defined vascular-associated neuroimmune niches, positioning NK cells within specialized microenvironments that support IL-17-dependent repair.

## Cholinergic modulation promotes repair across species

Having established that localized cholinergic modulation promotes diabetic wound repair through a CHRM1-dependent, IL-17-associated neuroimmune program in mice, we next asked whether this reparative response extends across clinically relevant large animal and human skin models. Because no single experimental system fully recapitulates the complexity of DFUs, therapeutic efficacy was evaluated across complementary *in vivo* and *ex vivo* models.

**Fig. 1.**
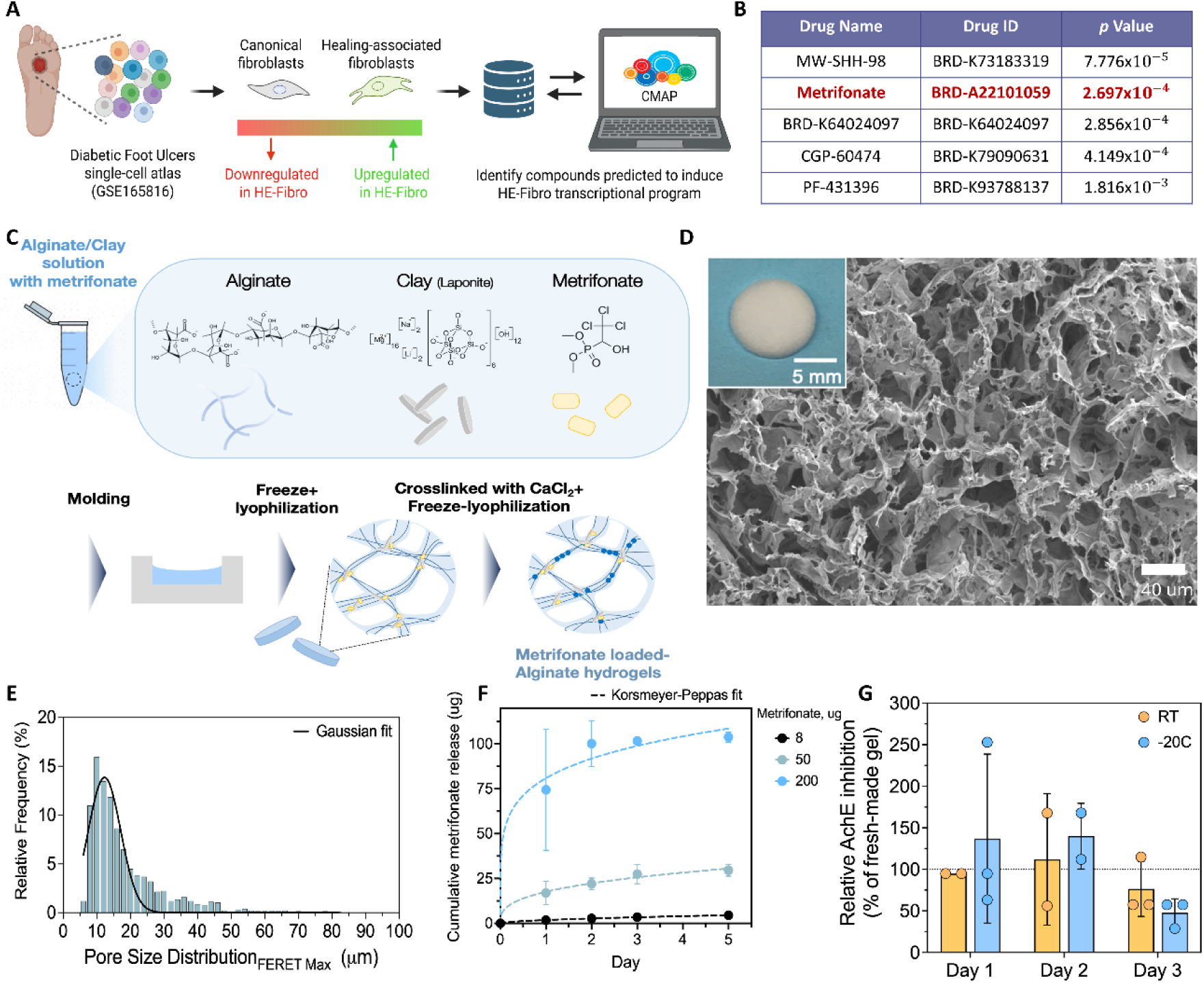
Computational identification of metrifonate and development of a sustained-release hydrogel for localized cholinergic modulation. **A**, Computational drug repurposing workflow identifying compounds predicted to induce the healing-associated fibroblast transcriptional program derived from the human diabetic foot ulcer atlas. B, Top-ranked candidate compounds identified by Connectivity Map (CMAP) analysis are shown. *p*-values were determined using Kolmogorov-Smirnov enrichment scoring evaluated against random gene-set permutation distributions. C, Schematic illustrating fabrication of metrifonate-loaded alginate-clay hydrogels (Met-gel). D, Scanning electron micrograph and representative macroscopic image (inset) of the hydrogel. E, Hydrogel pore size distribution. F, Cumulative (low, medium and high dose) metrifonate release profiles from alginate-clay hydrogels. G, Relative acetylcholinesterase (AchE) inhibition activity of metrifonate released from metrifonate-loaded alginate/clay hydrogels stored at RT and -20 °C for 17 days, compared to freshly made hydrogels. The AChE inhibition activity of freshly made hydrogels is represented by a dotted line at 100%. Data are mean ± s.d. (*n* = 2-3 independent hydrogels per condition).

**Fig. 2.**
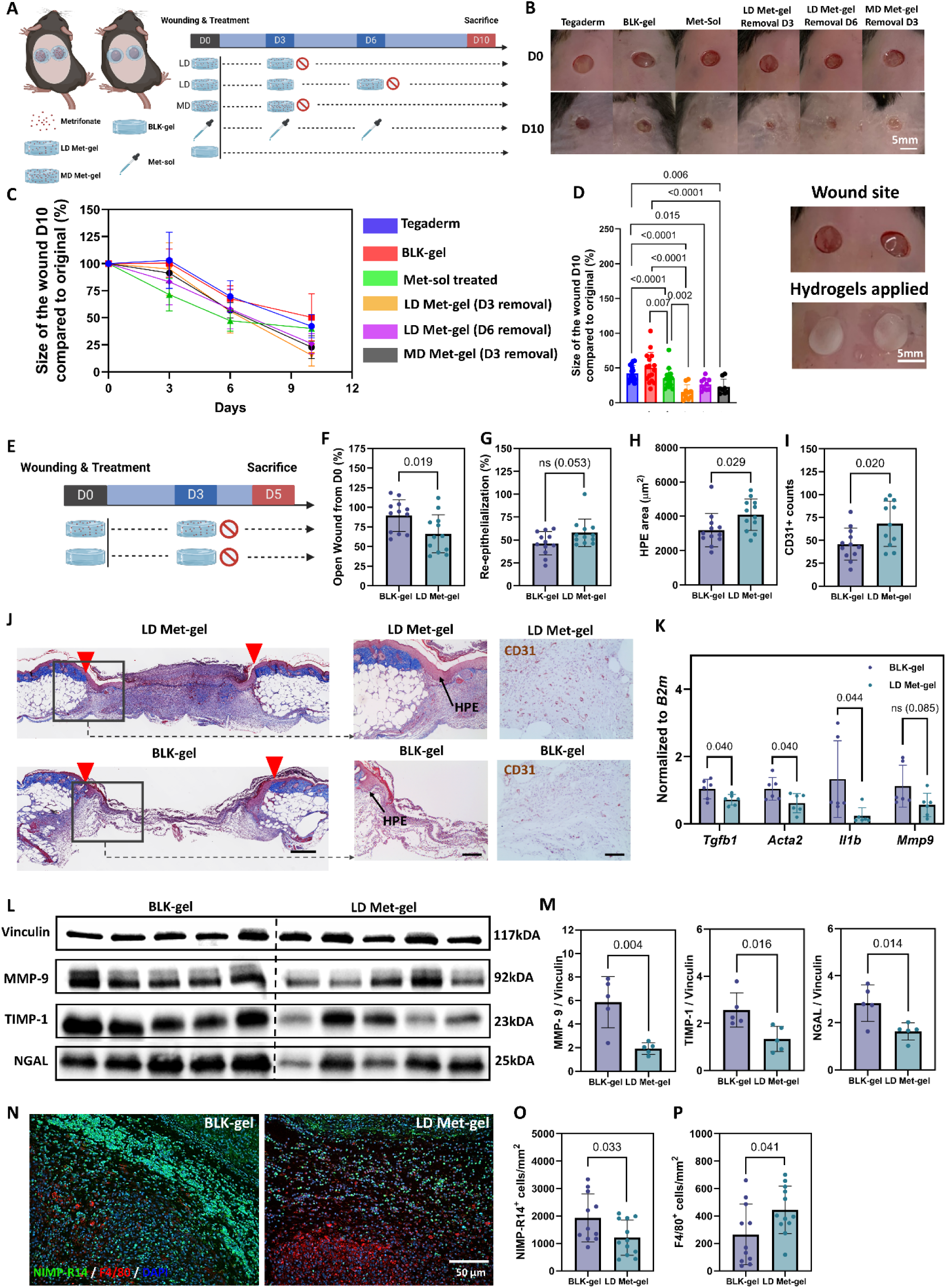
Localized sustained delivery of metrifonate accelerates diabetic wound repair. **A,** Experimental design comparing topical soluble metrifonate (Met-sol), low-dose metrifonate hydrogel (LD Met-gel) removed on day (D) 3 or D6, medium-dose metrifonate hydrogel (MD Met-gel) removed on D3, Tegaderm dressing, and blank hydrogel (BLK-gel) controls in *db/db* mice. **B-D,** Representative wound images **(B)**, wound closure over time **(C)**, and wound closure at day 10 **(D)** following treatment with the indicated formulations. Representative images of hydrogel application after wounding (**D**, far right). Scale bars = 5 mm. **E,** Experimental design for length of treatment and tissue collection time points following application of BLK-gel or LD Met-gel. **F-I,** Quantification of wound closure **(F)**, re-epithelialization **(G)**, hyperproliferative epidermis (HPE) area **(H)**, and CD31⁺ vessel density **(I)**. **J,** Representative histological sections stained with Masson’s trichrome (MTS) and immunohistochemical staining for CD31. Red arrowheads indicate wound margins. Black arrows point to the HPE. Scale bars, 100 μm. **K,** RT-qPCR analysis of wound repair-associated genes on day 5. **L,M,** Representative immunoblots **(L)** and quantification **(M)** of MMP-9 (left), TIMP-1 (center) and NGAL (right) protein expression. Vinculin was used as loading control. **N-P,** Representative immunofluorescence images of neutrophils (NIMP-R14⁺ cells; green) and macrophages (F4/80⁺ cells; red) **(N)**, with quantification of NIMP-R14⁺ cells **(O)** and F4/80⁺ cells **(P)**. DAPI was used for nuclear counterstain. Scale bar, 50 μm. Data are mean ± s.d. Sample sizes are indicated in the figure. Statistical analyses were performed using mixed-effects models or two-sided unpaired *t*-tests as appropriate.

**Fig. 3.**
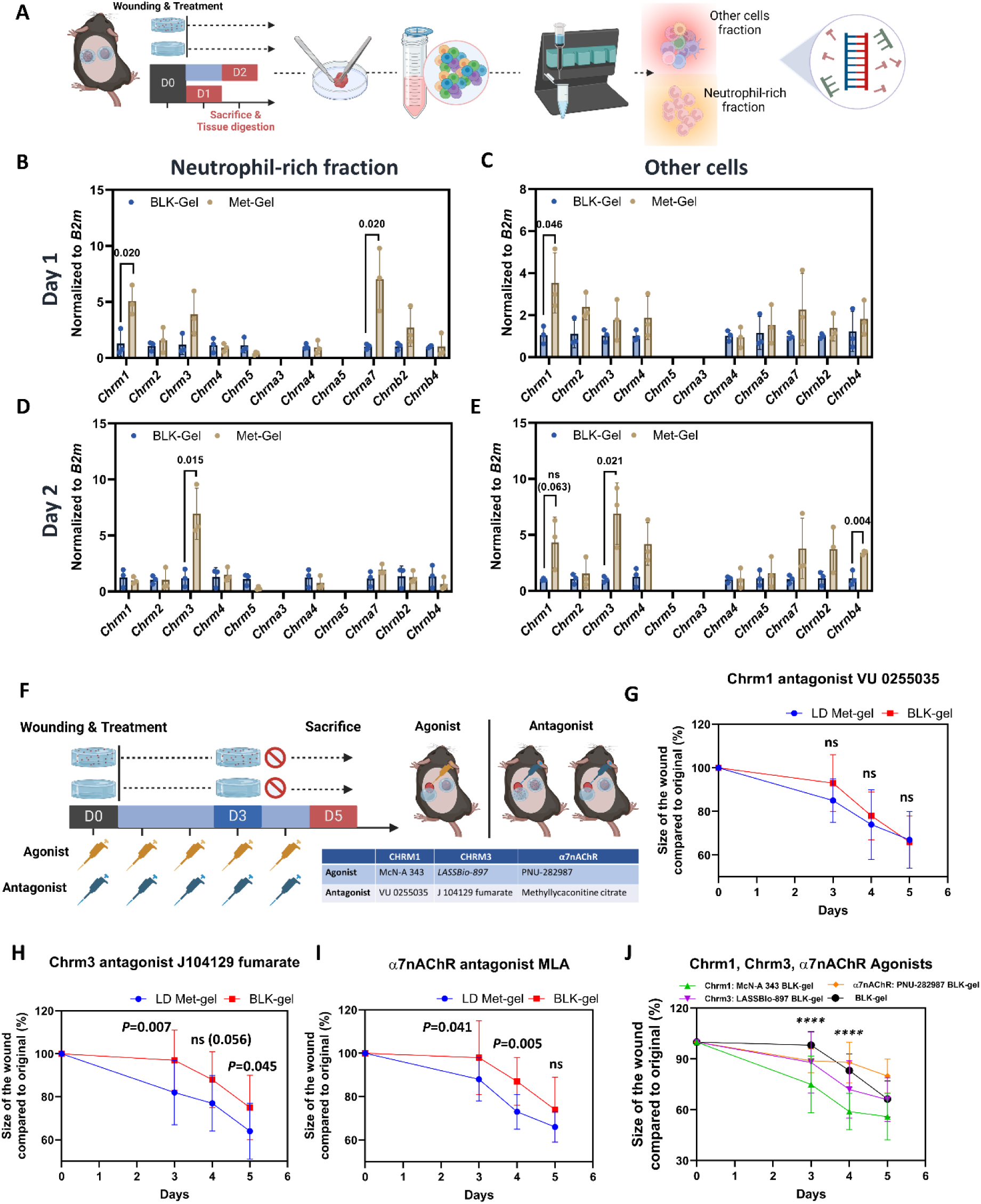
Acetylcholinesterase inhibition selectively signals through CHRM1 during the early inflammatory phase of diabetic wound repair. **A,** Experimental design for isolation of neutrophil-enriched and non-neutrophil wound cell fractions from day 1 and day 2 wounds following treatment with low-dose metrifonate hydrogel (LD Met-gel) or blank hydrogel (BLK-gel). Wounds were enzymatically dissociated and neutrophils were isolated by magnetic-activated cell sorting (MACS) prior to downstream molecular analyses. **B-E,** RT-qPCR analysis of muscarinic and nicotinic acetylcholine receptor expression in neutrophil-enriched **(B,D)** and non-neutrophil **(C,E)** fractions isolated from day 1 **(B,C)** and day 2 **(D,E)** wounds (*n*=3 per group). **F,** Experimental design for pharmacological interrogation of cholinergic receptor signaling using selective muscarinic and nicotinic receptor agonists and antagonists. **G-I,** Wound closure following topical treatment with LD Met-gel or BLK-gel in the presence of selective antagonists targeting CHRM1 **(G)**, CHRM3 **(H)**, or α7 nicotinic acetylcholine receptors **(I)**. Data are mean ± s.d. (*n* = 12). **J,** Wound closure following topical administration of selective agonists targeting CHRM1, CHRM3, or α7 nicotinic acetylcholine receptors in combination with BLK-gel. Data are mean ± s.d. (*n* = 4-12). **** (*P*<0.0001): Day 3 McN-A 343 BLK-gel vs. Day 3 BLK-gel, and Day 5 McN-A 343 BLK-gel vs. Day 5 BLK-gel. Statistical analyses were performed using two-sided unpaired *t*-tests or mixed-effects models with multiple-comparison correction, as appropriate.

**Fig. 4.**
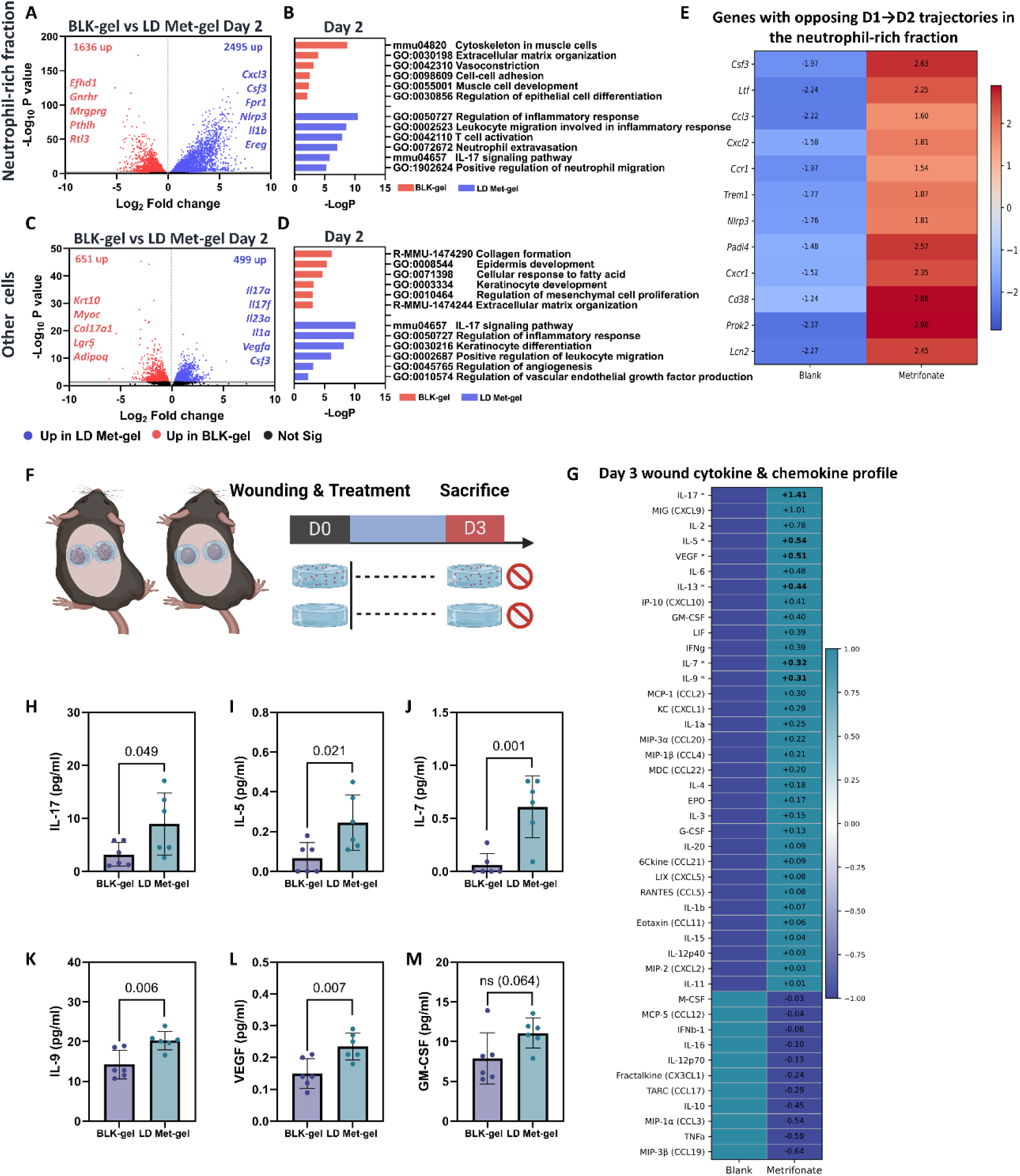
Early cholinergic modulation induces inflammatory and IL-17-associated transcriptional programs in diabetic wounds. **A,** Volcano plot showing differentially expressed genes (DEGs) in neutrophil-enriched fractions isolated from day 2 wounds treated with low-dose metrifonate hydrogel (LD Met-gel) or blank hydrogel (BLK-gel). **B,** Functional enrichment analysis of DEGs identified in neutrophil-enriched fractions on day 2. **C,** Volcano plot showing DEGs in non-neutrophil wound cell fractions isolated from day 2 wounds treated with LD Met-gel or BLK-gel. **D,** Functional enrichment analysis of DEGs identified in non-neutrophil wound cell fractions on day 2. **E,** Heatmap showing representative significantly DEGs with opposing temporal trajectories between blank gel-and metrifonate-treated wounds in the neutrophil-rich fraction. Values indicate log2 fold change in gene expression from day 1 to day 2 (D2 versus D1) within each treatment group. Negative values indicate higher expression at day 1 and positive values indicate higher expression at day 2. **F,** Experimental design for multiplex cytokine profiling of wound lysates collected on day 3 following treatment (*n*=6 wounds per treatment group). **G,** Heatmap showing the mean fluorescence intensity (MFI) of 44 cytokines, chemokines and growth factors measured by Luminex in day 3 wound tissue from blank gel- and metrifonate gel-treated mice (n = 6 per group). Values represent mean MFI across biological replicates and are row-scaled following log2 transformation to visualize treatment-associated changes within each analyte. Analytes are ordered by decreasing log2 fold change (metrifonate/blank), with log2 fold-change values indicated within the metrifonate column. Asterisks denote analytes significantly different between treatment groups (P < 0.05, two-tailed unpaired Student’s t-test). **H-M,** Quantification of IL-17 **(H)**, IL-5 **(I)**, IL-7 **(J)**, IL-9 **(K)**, VEGF **(L)** and GM-CSF **(M)** protein abundance in day 3 wound lysates. RNA sequencing was performed on neutrophil-enriched and non-neutrophil fractions isolated from *n* = 3 biologically independent wounds per treatment group. Cytokine measurements are presented as mean ± s.d. Statistical significance was determined using two-sided unpaired *t*-tests.

**Fig. 5.**
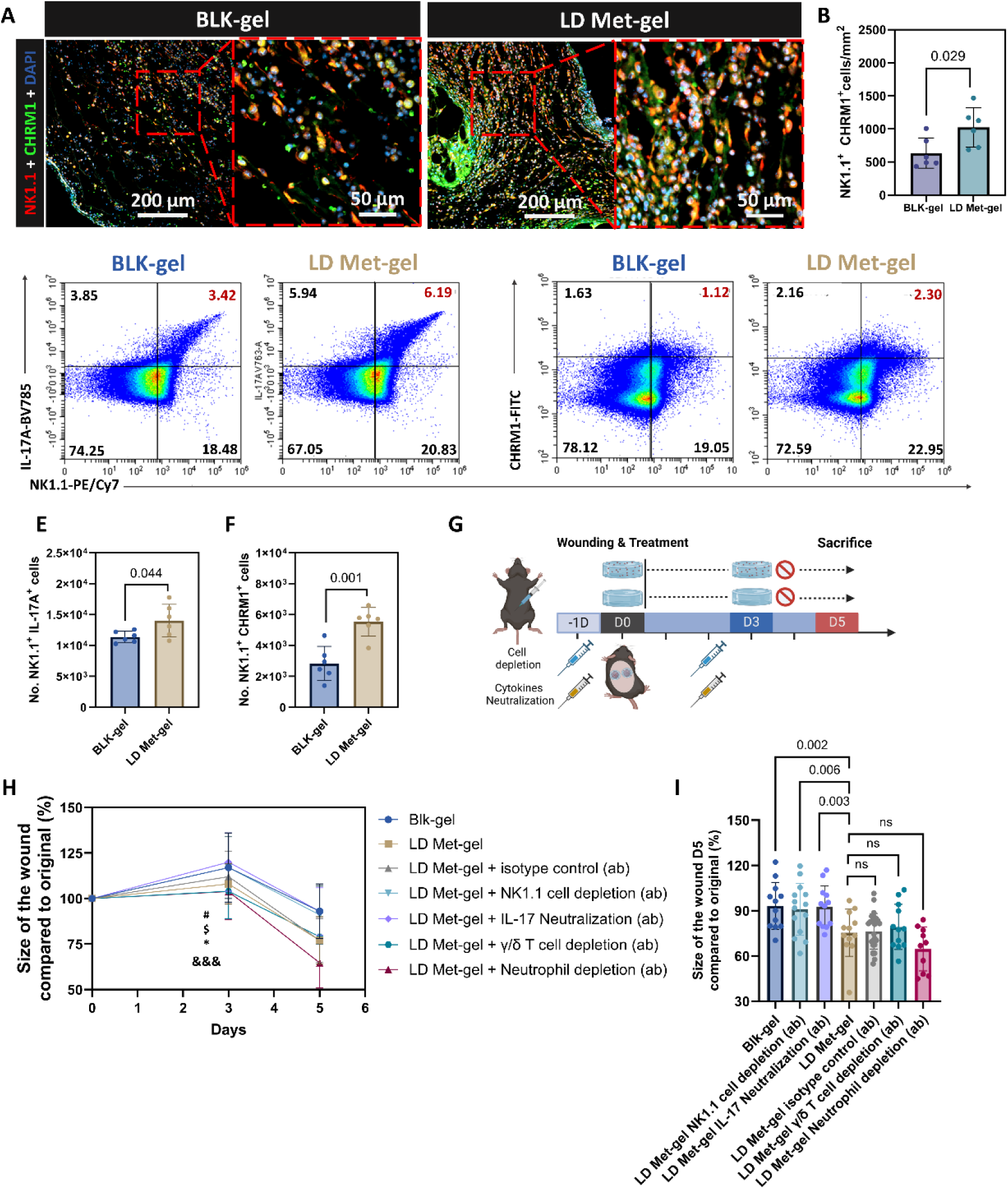
CHRM1-responsive NK1.1+ lymphocytes mediate IL-17A-dependent diabetic wound repair. **A,B,** Representative immunofluorescence images (A) and quantification (B) of CHRM1 expressing NK1.1⁺ cells in day 3 wounds treated with low-dose metrifonate hydrogel (LD Met-gel) or blank hydrogel (BLK-gel). **C-F,** Representative flow cytometry plots of IL-17A⁺NK1.1⁺ cells **(C)** and CHRM1⁺NK1.1⁺ cells **(D),** with corresponding quantification **(E,F)** of each population. Data are mean ± s.d. Sample sizes are indicated in the figure. Statistical significance was determined using two-sided unpaired *t*-tests, as appropriate. **G,** Experimental design for immune cell depletion and cytokine neutralization studies. **H,** Wound closure following depletion of NK1.1⁺ cells, γδ T cells, or neutrophils, or following IL-17A neutralization in mice treated with LD Met-gel or BLK-gel. Data are mean ± s.d. (*n* = 10-12). #: Day 3 LD Met-gel vs Day 3 BLK-gel; $ (P<0.05) LD Met-gel+γ/δ T cell depletion vs BLK-gel on Day 3; * (P<0.05) Day 3 LD Met-gel+Neutrophils depletions vs Day 3 BLK-gel; CCC (P<0.001) LD Met-gel+isotype control vs BLK-gel on Day 3**. I,** Size of open wound on Day 5. Data from the LD Met-gel isotype controls in H and J represent the combined data from the corresponding isotype control antibodies used for cell depletion or cytokine neutralization. *P* values were determined using mixed-effects model with Days and Treatment as fixed effects, followed by Fisher’s LSD multiple-comparison test.

**Fig. 6.**
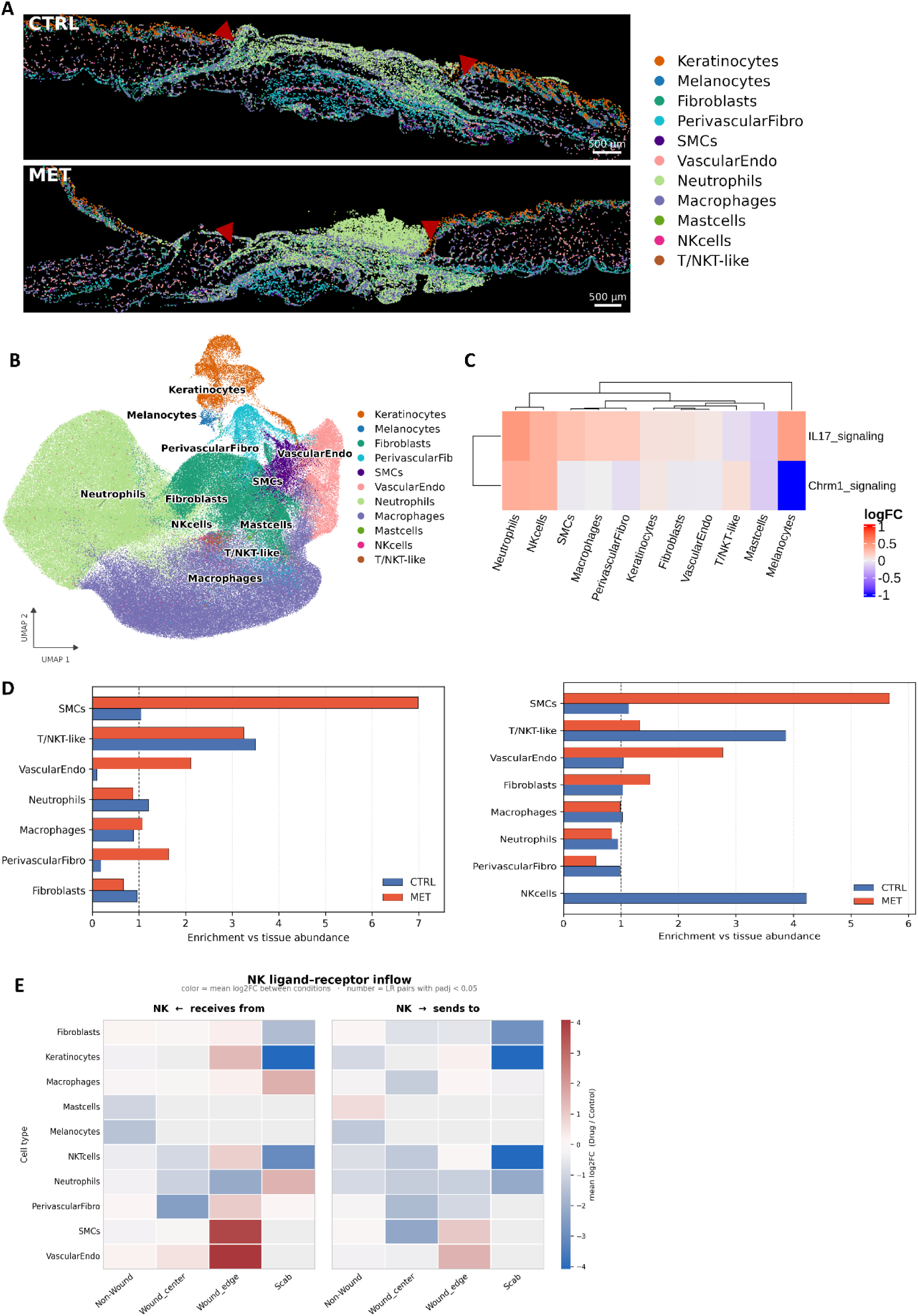
Spatial transcriptomics identifies vascular-associated neuroimmune niches induced by cholinergic modulation during diabetic wound repair. **A,** Representative Xenium spatial transcriptomic maps of day 3 diabetic wounds treated with low-dose metrifonate hydrogel (MET) or blank hydrogel (CTRL). Cells are colored according to annotated cell identity. Red arrowheads indicate wound margins. **B,** Uniform Manifold Approximation and Projection (UMAP) visualization of all cells identified by Xenium spatial transcriptomics following unsupervised clustering and cell-type annotation (*n* = 237,213 cells from 4 wounds per treatment group). **C,** Cell type-specific IL-17 signaling and CHRM1-associated pathway activity across annotated cell populations in LD Met-gel (MET) and BLK-gel wounds (CTRL). Heatmaps depict normalized pathway scores. **D,** Spatial neighborhood enrichment analysis showing preferential association of NK cells with smooth muscle cells (SMCs) and perivascular fibroblasts within the wound edge (left) and wound center (right). **E,** Ligand-receptor interaction analysis of NK cells showing predicted incoming and outgoing signaling interactions with neighboring cell populations. Scale bars, 500 μm.

We first assessed metrifonate in a diabetic Yucatan minipig model, which closely mirrors human skin architecture, immune composition, and wound healing dynamics (*40*). Diabetes was induced using alloxan, followed by creation of full thickness dorsal wounds after establishment of a sustained hyperglycemic state (*41*). Metrifonate loaded gel or blank gel was applied immediately following injury and removed after four days, emulating the localized sustained delivery strategy used in murine studies (**Fig. 7A**). Metrifonate treatment significantly improved wound closure across the 21-day observation period (general linear model, treatment effect, *P* = 0.015), with significantly greater closure evident at day 4 (*P* = 0.028; **Fig. 7B, C**). Histological analyses demonstrated enhanced re-epithelialization (**Fig. 7D**), increased epidermal thickness, and elevated angiogenesis in drug treated wounds (**fig. S10A-D**). By day 21, metrifonate treated wounds displayed reduced abundance of CD45⁺ immune cells, consistent with timely immune resolution and transition into tissue remodeling (**fig. S10E, F**). To assess early immune signaling, wound exudate recovered from the hydrogels during the initial post-injury period (Day 4) was analyzed by cytokine profiling (**table S3**). Metrifonate treatment resulted in increased IL-17A protein levels, mirroring findings in the murine model and supporting conservation of IL-17 associated immune activation (**Fig. 7E**). Additional cytokines IL-8 and IL-12, implicated in innate immune recruitment and activation were also elevated, consistent with coordinated early inflammatory engagement (**fig. S10G, H**) (*42, 43*). Gene expression analyses from Day 21 further revealed higher expression of growth factors associated with immune regulation, angiogenesis, and keratinocyte migration (**fig. S10I-K)** (*44*).

**Fig. 7.**
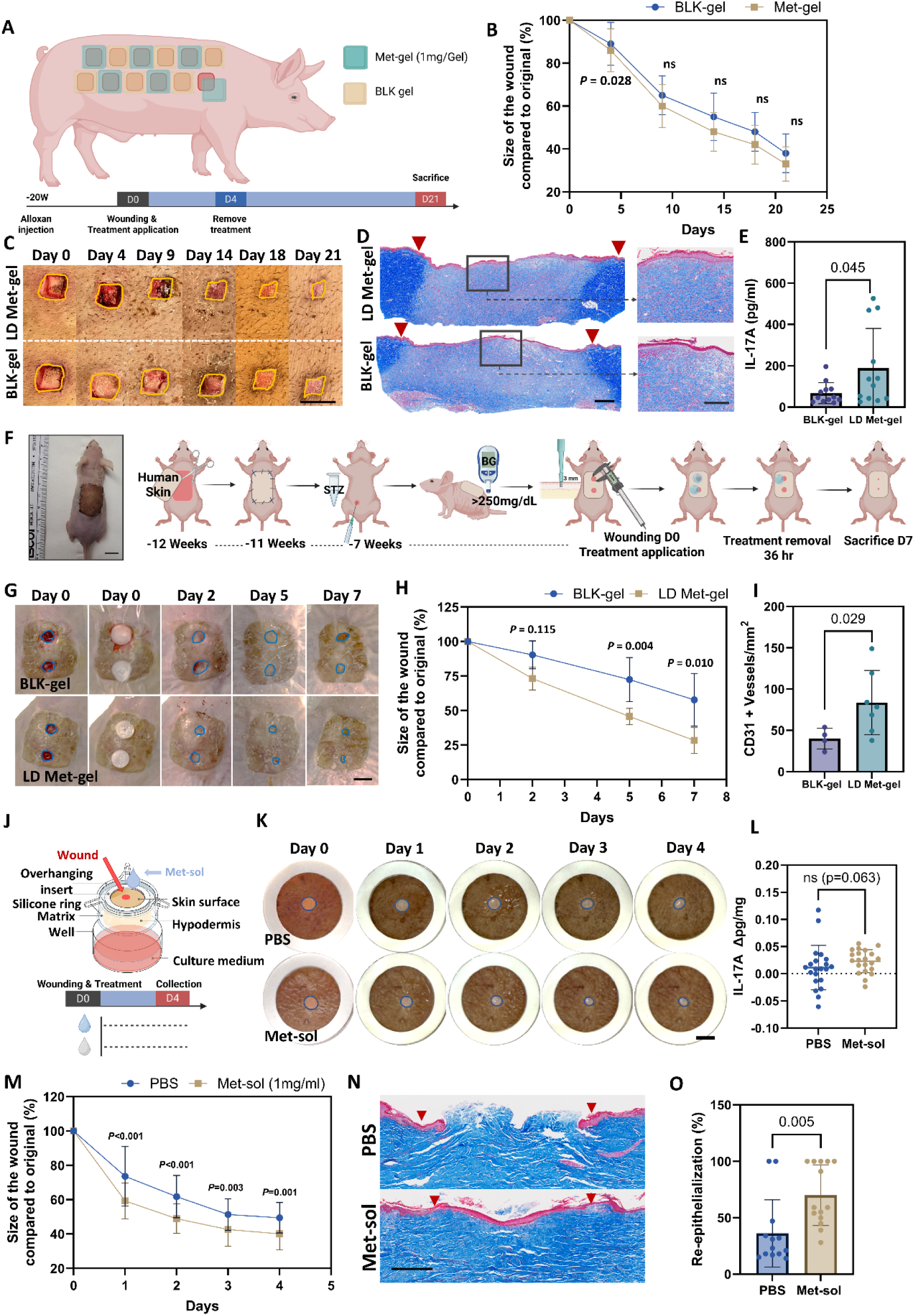
Localized cholinergic modulation promotes wound repair across clinically relevant translational models. **A,** Experimental design for evaluation of low-dose metrifonate hydrogel (LD Met-gel) in a diabetic porcine wound healing model. **B-D,** Wound closure over time. Data are mean ± s.d. (*n* = 12 wounds per treatment from one pig). *P* values were determined using a general linear model with day and treatment as fixed factors. Overall treatment effect, *P* = 0.015. *P* values shown at individual time points represent Fisher’s LSD comparisons between Met-gel and BLK-gel. **(B)**, representative macroscopic images with yellow lines demarcating wound edges **(C)**, and Masson’s trichrome-stained wound sections at day 21 **(D)**. Red arrowheads indicate wound margins. Scale bars, 1 cm **(C)** and 500 μm **(D)**. **E,** IL-17A concentration in wound exudates collected on day 4. **F,** Representative mouse image with transplanted skin (left) and experimental design (right) for the diabetic human skin xenograft model. **G-I,** Representative wound images with blue lines demarcating wound edges; scale bar= 5 mm **(G)**, wound closure over time. Data are mean ± s.d. (*n* = 4-7). General treatment effects: *P* = 0.02, determined by mixed effects model **(H)**, and quantification of CD31⁺ vessel density **(I)**. **J,** Experimental design for the ex vivo human skin wound model. The schematic illustration adapted from Genoskin Inc. **K-O,** Representative wound images with blue lines demarcating wound edges; scale bar= 5 mm **(K)**, change in IL-17A secretion between day 2 and day 4 **(L)**, wound closure over time, Data are mean ± s.d. (*n* = 20-21). General treatment effects: *P* = 0.001, determined by mixed effects model **(M)**, Masson’s trichrome-stained wound sections; red arrowheads denote wound margins; scale bar= 500 μm **(N)**, and re-epithelialization at day 4 **(O)**. Data are mean ± s.d. Sample sizes are indicated in the figure. Statistical analyses were performed using two-sided unpaired *t*-tests (**E**, **I**, and **O**), or Mann-Whitney tests (**I**), as appropriate. *P* values in **H** and **M** for individual comparisons, as determined by Fisher’s LSD multiple-comparisons test following a mixed-effects model with day and treatment as fixed effects, are shown in the figure.

To further extend translational relevance, we evaluated metrifonate in two complementary human skin models. First, we employed a model in which split thickness human skin was transplanted onto immunodeficient nude mice, followed by induction of diabetes (**Fig. 7F**). After establishment of stable hyperglycemia, wounds were generated onto the human grafts and treated with metrifonate-loaded or blank gel (**Fig. 7G**) (*45*). In agreement with murine and porcine findings, treatment significantly accelerated wound closure in human skin grafts (**Fig. 7H**), enhanced angiogenic responses, and resulted in greater numbers of Ki67+ proliferating cells in both the epidermal and dermal compartments of the wounds (**Fig.7I** and **fig. S10L-Ǫ)**.

We next assessed metrifonate in an *ex vivo* human skin wound model derived from surgical tissue obtained from multiple independent donors (**Fig. 7J**). In this system, wounds were treated with drug solution or vehicle control immediately after injury, and closure was quantified over time (**Fig. 7K**). Treatment resulted in enhanced wound edge advancement across donors (**Fig. 7M-O**). Cytokine analysis of conditioned media additionally revealed a trend toward increased IL-17A secretion following treatment (**Fig.7L**). Importantly, treated human wounds exhibited increased numbers of CHRM1⁺ and IL-17A⁺ cells (**fig. S10R-U**), recapitulating key cellular features of the cholinergic immune response identified in murine wounds.

Across murine, porcine, and human skin models, localized metrifonate treatment consistently accelerated wound healing, enhanced re-epithelialization and angiogenesis, and engaged IL-17-associated immune signaling despite substantial differences in tissue architecture and immune composition. The concordance of therapeutic responses across complementary experimental systems indicates that the reparative program identified in mice is not species restricted and supports the translational potential of topical cholinergic modulation for diabetic wound repair.

## Discussion

Our findings identify localized cholinergic modulation as a mechanism for restoring productive inflammation in diabetic wounds. Although metrifonate was identified from a healing-associated fibroblast transcriptional signature, its therapeutic activity was mediated through the immune microenvironment, revealing an unexpected connection between cholinergic signaling and tissue repair. AChE inhibition induced an early inflammatory program characterized by prominent IL-17 signaling, and IL-17A neutralization abolished therapeutic efficacy. Although excessive or persistent IL-17 signaling has been associated with chronic inflammation and impaired healing (*4C-48*), IL-17 can also support tissue repair in a context- and time-dependent manner (*4S-51*). Our findings suggest that defective diabetic wound healing may therefore reflect not only excessive inflammation, but also failure to appropriately initiate reparative acute-like inflammatory programs. The healing-associated fibroblast state that enabled identification of metrifonate may, in turn, represent a downstream manifestation of this permissive immune environment.

Non-neuronal cholinergic signaling is increasingly recognized as a regulator of immune function (*52*), but its contribution to wound repair remains poorly understood. Pharmacological interrogation identified CHRM1 as required for the therapeutic effects of metrifonate, while cellular analyses implicated CHRM1-expressing NK1.1+ innate lymphocytes in this response. Depletion of NK1.1+ cells abolished therapeutic efficacy, whereas depletion of γδ T cells or neutrophils did not. Spatial profiling further localized NK cells to vascular-associated microenvironments enriched for interactions with smooth muscle cells, perivascular fibroblasts and endothelial cells, providing a potential framework through which a restricted cholinergic-responsive immune population could coordinate broader tissue repair. Wound associated NK cells can regulate the balance between inflammation, angiogenesis and tissue repair, with their functional state shaped by local environmental cues (*53*). The precise identity of the required NK1.1+ population remains unresolved, however, and cell-specific genetic studies will be required to establish whether CHRM1 signaling within conventional NK cells directly drives the IL-17A-dependent response.

Therapeutic efficacy across diabetic mouse and porcine wounds, together with human skin models, suggests that the reparative effects of localized cholinergic modulation are not restricted to a single experimental system and species. Given that no single animal model fully recapitulates human chronic wound healing, efficacy across complementary models provides important translational support, particularly in porcine wounds, which more closely reproduce the architecture and repair characteristics of human skin (*54*). The use of an alginate-based delivery platform already established in wound care (*55*), together with the previous long history of clinical applications for AChE inhibitors (*5C*), further supports the potential for therapeutic repurposing and progression toward clinical evaluation per FDA guidelines. More broadly, our findings suggest that chronic wounds may benefit from strategies that restore the timing and organization of endogenous inflammatory responses rather than simply suppressing inflammation. By linking localized AChE inhibition to CHRM1-associated innate lymphocyte responses and IL-17A-dependent repair, this study identifies cholinergic signaling as a previously unrecognized regulatory axis in diabetic wound healing and provides a framework for therapeutically re-engaging productive inflammation in chronic wounds.

## Data availability

The bulk RNA-sequencing data generated in this study have been deposited in the Gene Expression Omnibus (GEO) under accession number GSE335615. The Xenium spatial transcriptomic data generated in this study have been deposited in Zenodo under DOI: 10.5281/zenodo.20550575. The previously published single-cell RNA-sequencing dataset of human diabetic foot ulcers used to derive the healing-associated fibroblast transcriptional signature is publicly available under accession number GSE165816. Source data underlying the figures are provided with this paper.

## Acknowledgements

We thank the Beth Israel Deaconess Medical Center Flow Cytometry Core Facility for technical support and the NIH Tetramer Core Facility for providing reagents. Schematics were created with BioRender. We thank Daniel Hoft for providing Mr1-deficient mice and Richard Blumberg for providing Jα18-deficient mice. This work was supported by NIH grants R44-DK141401 (A.V. and D.J.M.) and K01-DK139434 (G.T.), a Wound Healing Foundation 3M Fellowship (G.T.), a Pilot and Feasibility Award from the Joslin Diabetes Center (P30-DK036836) (G.T.), and Massachusetts Life Sciences Center (grant 26018) (G.T.). Z.L. was supported by a Moderna Global Fellowship (award no. 2022-26). M.G.F. was supported by a São Paulo Research Foundation (FAPESP) Research Internship Abroad (BEPE) fellowship (grant no. 2024/18496-7). N.L-G. was supported by the CRIS Cancer Foundation (Outback2022_14, co-financed by Fundación ADEY). L.M.A. thanks the Basque Government, Spain, for awarding the Ph. D. grant (PRE_2022_1_0053). A.V. received support from the National Rongxiang Xu Foundation.

## Author contributions

G.T., D.J.M. and A.V. conceived and designed the study. Z.L. performed the mouse experiments, flow cytometry, RT-qPCR, western blotting and ELISA experiments. G.T. performed flow cytometry experiments and designed the Xenium gene panel. S.L. developed and characterized the metrifonate-loaded hydrogel, with assistance from N.D.; N.L.-G. and L.M.A. contributed to hydrogel fabrication. N.K. performed the healing-associated fibroblast and Connectivity Map analyses that led to the identification of metrifonate, under the supervision of I.S.V. G.T. performed downstream analysis of the bulk RNA-sequencing data. S.W. and A.A. performed the Xenium spatial transcriptomic experiments, and S.U.N. performed the spatial transcriptomic analyses under the supervision of I.S.V. E.A.V, H.N., I.M., M.G. and G.C. assisted with mouse experiments. Z.L., M.C., E.A.V., H.N., E.W., M.G., G.C., J.H. and J.G. performed the porcine wound healing studies. Z.L., I.M., E.W. and B.J.S. performed the human skin xenograft studies. Z.L. and M.G.F. performed the ex vivo human skin studies. Z.L., E.A.V., M.G., G.C., J.H. and M.G.F. performed immunostaining, imaging and quantitative image analyses. Z.L., G.T. and S.L. generated the figures and wrote the manuscript. G.T., D.J.M. and A.V. acquired funding. D.J.M. and A.V. supervised the study. All authors reviewed and edited the manuscript.

## Supplementary Materials for

**This PDF file includes:**

Materials and Methods

Figs. S1 to S10

Tables S1 to S7

**Other Supplementary Materials for this manuscript include:**

Data S1 to S3

### Materials and Methods

#### ASGARD Connectivity Map drug-repurposing analysis

A transcriptional profile representing the healing-associated fibroblast (HE-Fibro) state was derived from our previously published single-cell analysis of human DFUs (GSE165816). Foot-skin cells annotated by SCINA(*57*) as HE-Fibro or canonical fibroblasts were analyzed, retaining genes detected in at least 10% of cells in either population. Differential expression was assessed using limma(*58*), with patient and study center included as covariates. The resulting HE-Fibro versus fibroblast differential expression profile was ranked by log2 fold change and queried against Connectivity Map/LINCS L1000 perturbational profiles (GSE70138 and GSE92742) using ASGARD(*19*) with positive connectivity to identify compounds predicted to induce the HE-Fibro transcriptional state. Among 11,908 perturbagens evaluated, metrifonate (BRD-A22101059) ranked second (nominal *P* = 2.70 × 10⁻⁴).

#### Fabrication of metrifonate-incorporated alginate gels

Metrifonate-incorporated alginate gels animal studies were fabricated. The general process and alterations for specific studies are described below. A 39 mg/mL alginate solution (PRONOVA UP MVG, Sigma-Aldrich, 42000101), a 20.6 mg/mL laponite solution (BYK USA Inc.), and a 50 mg/mL metrifonate stock solution (Trichlorfon, Supleco, 45698) were prepared separately in deionized (DI) water. The metrifonate and laponite solutions were mixed for 30 minutes and subsequently combined with the alginate solution to achieve final concentrations of 10 mg/mL laponite, and 20 mg/mL alginate, with a total metrifonate payload of 8 µg (for rodent studies) or 1,000 µg and 5,000 µg (for porcine studies) per gel. The mixture was cast into molds (1 cm diameter circular molds for the *db/db*, WT and knockout mouse studies, 5 mm diameter for nude mice study, and 2 cm x 2 cm square molds for the porcine study), frozen at -20 °C, and lyophilized. After lyophilization, the gels were crosslinked by incubation in 0.1 M CaCl_2_ (for rodent studies) or 0.02 M CaCl_2_ (for porcine studies) for 30 minutes on an orbital shaker. The crosslinked gels were washed three times with DI water, frozen at -20 °C, and lyophilized again. The final dried gels were stored at -20 °C and rehydrated with PBS immediately prior to wound application.

#### Scanning electron microscopy (SEM) characterization of metrifonate-incorporated alginate gels

Metrifonate-loaded alginate gels were lyophilized for 24 h and coated with a 10 nm layer of Pt/Pd using an EMS 150T ES Sputter/Carbon Coater (Electron Microscopy Sciences, USA). The samples were imaged using a Zeiss Gemini 360 field-emission scanning electron microscope (FE-SEM, Zeiss, Germany) operated at 7 kV. The SEM images were analyzed using ImageJ software (NIH, USA). To accurately identify the pore structure, the Trainable Weka Segmentation plugin was utilized. Following segmentation, the pore size distribution was determined based on the maximum Feret’s diameter. For statistical quantification, a minimum of 300 pores were measured per sample (*n*=3).

##### Metrifonate Release Studies

Metrifonate-incorporated alginate/laponite gels were incubated in PBS (pH 7.4) at 37 °C under constant shaking. At each time point, an aliquot of the supernatant was collected for release assays and replaced with an equal volume of PBS. An Acetylcholinesterase inhibitor screening kit (Sigma-Aldrich, MAK324) was used to quantify the inhibitory activity of the released metrifonate, respectively. Metrifonate inhibitory activity in the sample was measured spectrophotometrically at 412 nm, a yellow product from 5,5’-dithiobis (2-nitrobenzoic acid) (DTNB) and calculated using a calibration curve generated from standard solutions.

##### Mathematical modeling of drug release kinetics

To elucidate the release mechanism and kinetics of metrifonate from the alginate/nanoclay hydrogels, the in vitro drug release data were fitted to the Korsmeyer-Peppas (K-P) model and Peppas-Sahlin (P-S) mathematical models. The K-P model is expressed as *M*_*t* ⁄ *M*_∞ = *kt*^*n*, where *M*_*t* ⁄ *M*_∞ is the fraction of drug released at time t, k is the release rate constant, and n is the release rate exponent (diffusional exponent). The P-S model is expressed as *M*_*t* ⁄ *M*_∞ = *k*_1 *t*^*m* + *k*_2 *t*^2*m*, where *k*_1represents the Fickian diffusional constant, *k*_2 is the relaxation rate constant, and m is the Fickian diffusional exponent. To investigate the effect of drug loading on the release kinetics, the K-P model parameters (k and n) were plotted against the metrifonate-to-nanoclay ratios. The relationship between either the release rate constant (k) and the metrifonate-to-nanoclay ratio was modeled using a Hill equation to evaluate the saturation of the alginate/nanoclay binding capacity. Meanwhile, the correlation with the release rate exponent (n) was modeled using a power-law equation. All mathematical modeling and non-linear regression analysis were performed using GraphPad Prism 11.0 (GraphPad Software, USA).

##### In Vitro Swelling and Degradation studies

The swelling kinetics of gels were evaluated in PBS (pH 7.4). After recording the initial dry weight (Wd), samples were fully immersed in 1mL of PBS and incubated in sealed containers at 37 °C. At predetermined intervals, samples were removed, gently blotted with absorbent paper to remove surface buffer, weighed to determine wet weight (Ww), and immediately returned to the solution. The swelling ratio (g/g) was calculated using the following equation:

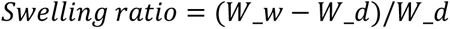

The macroscopic swelling kinetics of the hydrogels were modeled using a Voigt-based viscoelastic model(*59, 60*). The swelling data were fitted to the following equation:

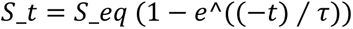

Where St is the swelling ratio at time t, Seq is the theoretical equilibrium swelling ratio, and *τ* is the relaxation time of the polymer network.

The degradation profiles of the gels were evaluated in PBS, pH 7.4. Lyophilized samples were first fully rehydrated to their maximum swelling capacity, and their initial wet weight (W0) was recorded. The samples were then incubated under the same conditions used for the swelling experiments. At predetermined intervals, weighed to record the degraded wet weight (Wt). The mass loss (%) was calculated as ((*WW*0 − *WWt*)/*WW*0) *x* 100.

##### FTIR analysis

The chemical composition of the hydrogels was analyzed using an attenuated total reflectance Fourier transform infrared (ATR-FTIR) spectrometer (Nicolet iS50 FTIR Spectrometer, Thermo Fisher Scientific) to verify the incorporation of clay nanoparticles (Laponite) within alginate gels. The pristine alginate and Laponite-loaded alginate hydrogels were lyophilized prior to analysis. The infrared absorption spectra of the samples were obtained. All spectra were recorded at room temperature in the wavenumber range of 1200 to 350 cm^-1^ with a resolution of 4 cm^-1^ and an accumulation of 32 scans per sample.

#### Animals

All animal experiments were performed in accordance with protocols approved by the Institutional Animal Care and Use Committee (IACUC) at Beth Israel Deaconess Medical Center (BIDMC). Male *db/db* mice (BKS.Cg-*Dock7m* +/+ *Leprdb*/J; strain no. 000642), male C57BL/6J mice (strain no. 000664), and male athymic nude mice (NU/J; *Foxn1nu*/*Foxn1nu*; strain no. 002019) were obtained from The Jackson Laboratory (Bar Harbor, ME, USA). MR1-deficient (*Mr1*−/−) mice on a C57BL/6 background were kindly provided by Daniel Hoft (Saint Louis University), and Jα18-deficient (*Traj18*−/−) mice (JAX strain no. 030524) were kindly provided by Richard Blumberg (Brigham and Women’s Hospital). Both strains were subsequently bred at BIDMC. A diabetic neutered male Yucatan miniature swine was obtained from Sinclair Bio Resources (Auxvasse, MO, USA).

#### *In vivo* diabetic mouse skin wound healing study

Male *db/db* mice were acclimated to the animal facility for at least 7 days before surgery. Hyperglycemia (blood glucose >250 mg dl⁻¹) was confirmed using a commercially available glucometer (Contour, Bayer). At 16 weeks of age, mice were anaesthetized with isoflurane, and the dorsal skin was shaved and depilated. Two 6-mm circular full-thickness excisional wounds were created on the dorsum using a biopsy punch (Integra Miltex), as previously described(*22*). Blank hydrogels (BLK-gel) or metrifonate-loaded hydrogels (Met-gel) were applied immediately after wounding, and the wounds were covered with an occlusive dressing (Tegaderm, 3M) to secure the hydrogels in place. Two Met-gel doses were evaluated: a low-dose formulation containing 8 μg metrifonate per wound and a medium-dose formulation containing 50 μg metrifonate per wound. Low-dose Met-gels were removed on either day 3 or day 6 after wounding, whereas medium-dose Met-gels were removed on day 3. Corresponding BLK-gels were removed at the same time points. Mice were monitored daily after surgery. Wounds were photographed and two perpendicular wound diameters were measured using digital calipers (Thermo Fisher Scientific, 14-648-17) on days 0, 3, 6 and 10. Wound area was calculated assuming an elliptical shape (A=πab), where a and b represent the semi-major and semi-minor axes, respectively, and expressed as the percentage of the initial wound area remaining relative to day 0. Mice were euthanized by CO₂ inhalation on days 1, 2, 3, 5 or 10 after wounding, depending on the experimental endpoint, and wound tissues were collected for downstream analyses.

#### *In vivo* evaluation of clinically used AChE inhibitors

Male *db/db* mice were treated topically with clinically used AChE inhibitors. Two 6-mm circular full-thickness excisional wounds were created on the dorsum as described above. Immediately after wounding, metrifonate (trichlorfon; Supelco, 45698), rivastigmine tartrate (Sigma-Aldrich, SML0881), donepezil hydrochloride (Sigma-Aldrich, D6821), or galantamine hydrobromide (Sigma-Aldrich, G1660) solution (50 μl at 1 mg ml⁻¹) or PBS vehicle control was applied directly to the wound area. AChE inhibitors were applied once daily from Day 0 through Day 3. Wounds were photographed and measured using digital calipers on Days 0, 3, 6, and 10. Mice were euthanized on Day 10. Wound size was measured as described above and expressed as the percentage of the initial wound area remaining relative to Day 0.

#### *In vivo* cell depletion and cytokine neutralization

Male *db/db* mice underwent immune cell depletion or cytokine neutralization before and/or during wound healing. For γδ T-cell or NK1.1⁺ cell depletion, mice received intraperitoneal injections of 400 μg anti-TCRγδ antibody (Bio X Cell, clone UC7-13D5, BE0070) or 400 μg anti-NK1.1 antibody (Bio X Cell, clone PK136, BE0036), respectively, 24 h before wounding, followed by 200 μg of the corresponding antibody on day 2 after wounding, as previously described(*61–63*). For neutrophil depletion, mice received 100 μg anti-Ly6G antibody (Bio X Cell, clone 1A8, BP0075) intraperitoneally 24 h before wounding, as previously described(*64*). Corresponding isotype control antibodies were Armenian hamster IgG (Bio X Cell, BE0091), mouse IgG2a (clone C1.18.4, BE0085) and rat IgG2a (clone 2A3, BP0089), respectively. For IL-17A neutralization, mice received 200 μg anti-IL-17A antibody (Bio X Cell, clone 17F3, BE0173) or mouse IgG1 isotype control antibody (clone MOPC-21, BE0083) intraperitoneally 24 h before wounding and again on day 2 after wounding, as previously described(*65*). Wounds were photographed and measured using digital calipers on days 0, 3 and 5. Mice were euthanized on day 5, and wound area was calculated as described above.

#### *In vivo* cholinergic receptor agonist and antagonist treatment

Male *db/db* mice were treated topically with selective cholinergic receptor agonists or antagonists following wounding. The CHRM1 agonist McN-A-343 (Tocris), CHRM3 agonist LASSBio-897 (BenChem), and α7 nicotinic acetylcholine receptor (α7nAChR) agonist PNU-282987 (Tocris) were administered individually at 50 μg per wound in 25 μl PBS before application of BLK-gel. For combined agonist treatment, the three agonists were administered together at a total dose of 50 μg per wound in 25 μl PBS. For receptor inhibition, the CHRM1 antagonist VU0255035 (Tocris), CHRM3 antagonist J104129 fumarate (Tocris), or α7nAChR antagonist methyllycaconitine citrate (Tocris) was applied topically at 50 μg per wound in 25 μl PBS before application of BLK-gel or low-dose Met-gel. Agonists and antagonists were administered once daily (Days 0 through 4) until the experimental endpoint on day 5. Wounds were photographed and measured as described above.

#### *In vivo* wound healing in STZ-induced diabetic WT, *Mr1*−/− and *Traj18*−/− mice

Male C57BL/6J wild-type (WT), *Mr1*−/− and *Traj18*−/− mice were used to assess the contribution of MAIT and iNKT cells to metrifonate-mediated wound repair. The origins and genetic backgrounds of the knockout strains are described above. Diabetes was induced beginning at 10 weeks of age in WT mice and between 10 and 21 weeks of age in *Mr1*−/− and *Traj18*−/− mice, depending on availability from in-house breeding. Mice received intraperitoneal injections of streptozotocin (STZ; 55 mg kg⁻¹) on four consecutive days, followed by a single 100 mg kg⁻¹ dose. Mice that did not develop hyperglycemia following this regimen received an additional 100 mg kg⁻¹ dose of STZ. Blood glucose was monitored using a glucometer, and mice with blood glucose >250 mg dl⁻¹ were considered diabetic. Hyperglycemia was maintained for at least 6 weeks before wounding. Mice were anesthetized with isoflurane, and two 6-mm full-thickness excisional wounds were created on the shaved and depilated dorsum using a sterile biopsy punch (Integra Miltex). BLK-gel or low-dose Met-gel containing 8 μg metrifonate per wound was applied immediately after wounding, and wounds were covered with an occlusive dressing (Tegaderm, 3M) to secure the hydrogels in place. Mice were monitored daily after surgery. Wounds were photographed and two perpendicular wound diameters were measured using digital calipers (Thermo Fisher Scientific, 14-648-17) on days 0, 3 and 5. Wound area was calculated assuming an elliptical shape as described above and expressed as the percentage of the initial wound area remaining relative to day 0. Mice were euthanized by CO₂ inhalation on day 5.

#### *In vivo* diabetic porcine skin wound healing study

A 5-month-old male Yucatan miniature swine was rendered diabetic by Sinclair Bio Resources by intravenous administration of alloxan monohydrate (100 mg kg⁻¹). Insulin glargine was administered daily to maintain fasting blood glucose concentrations of 300–500 mg dl⁻¹, and diabetes was maintained for approximately 5 months before wounding. General anaesthesia was induced by intramuscular administration of Telazol (10 mg kg⁻¹), followed by endotracheal intubation and maintenance with 1.5-3% inhaled isoflurane. Twenty-four full thickness excisional wounds (12 on each side of the dorsum), measuring 1.5 × 1.5 cm and approximately 0.6 cm in depth, were outlined using a sterile acrylic template to standardize wound placement and spacing. Wounds were separated by 2.5 cm and created using a no. 11 scalpel, forceps and surgical scissors. Twelve wounds were treated with BLK-gel and twelve with Met-gel containing 1 mg metrifonate per wound, with six wounds from each treatment group distributed on each side of the animal. Hydrogels were prepared as 2 × 2cm dressings, applied directly to the wounds immediately after wounding, and removed on day 4. Wounds were covered with Tegaderm (3M), and the surgical area was further protected using a custom fenestrated jacket (Lomir Biomedical). Postoperative analgesia, including buprenorphine and transdermal fentanyl, was administered according to institutional veterinary protocols. Wounds were photographed and measured using digital calipers (Thermo Fisher Scientific, 14-648-17) on days 0, 4, 9 and 14, 18 under anesthesia and on day 21 after euthanasia. Wound area was calculated from two perpendicular dimensions and expressed relative to the initial wound area on day 0. The animal was euthanized on day 21, and wound tissues were collected for downstream analyses.

#### *In vivo* diabetic human skin xenograft wound healing study

Six-week-old male athymic nude mice (*Foxn1*nu/*Foxn1*nu) were used as recipients for human skin xenografts. De-identified discarded split-thickness (700-800 μm) human abdominal skin from a 33-year-old female donor undergoing abdominoplasty was obtained from BioIVT (Westbury, NY, USA). Human skin transplantation and diabetes induction were performed as previously described(*45, 66*). Briefly, a 2 × 3 cm area of dorsal mouse skin was excised and replaced with a size-matched human skin graft, which was secured by suturing. Grafts were allowed to integrate for 5 weeks before diabetes induction. Diabetes was induced by four consecutive daily intraperitoneal injections of streptozotocin (STZ; 55 mg kg⁻¹), followed by a single 100 mg kg⁻¹ dose. Mice that did not develop hyperglycemia following this regimen received an additional 100 mg kg⁻¹ dose of STZ. Mice with blood glucose >250 mg dl⁻¹ were considered diabetic, and hyperglycemia was maintained for at least 6 weeks before wounding. Two 3-mm full-thickness excisional wounds were created within the human skin graft of each mouse. BLK-gel or Met-gel containing 8 μg drug per wound was applied immediately after wounding, and wounds were covered with Tegaderm (3M). Hydrogels were removed 36 h after application. Mice were monitored daily for signs of pain, distress or infection. Wounds were photographed and measured using digital calipers on days 0, 2, 5 and 7. Wound area was calculated from two perpendicular diameters assuming an elliptical shape, as described above, and expressed as the percentage of the initial wound area remaining relative to day 0. Mice were euthanized on day 7, and xenografted wound tissues were collected for downstream analyses.

#### Histology, immunohistochemistry and immunofluorescence

Mouse, human and porcine wound tissues were bisected through the wound center at the time of collection. One half was fixed in 10% neutral-buffered formalin and paraffin embedded, whereas the remaining half was snap frozen in liquid nitrogen and stored at -80 °C for subsequent RNA or protein extraction. Hematoxylin and eosin (H&E) and Masson’s trichrome staining were performed by the BIDMC Histology Core. Whole slide images were acquired using a Motic EasyScan Infinity slide scanner (iHisto, Salem MA). Re-epithelialization was quantified from H&E or Masson’s trichrome stained sections as the proportion of the wound bed covered by newly formed epithelium on QuPath v0.6.0.

For immunohistochemistry, paraffin-embedded sections were deparaffinized, rehydrated and subjected to antigen retrieval in citrate buffer (pH 6.0) using a pressure cooker for 15 min. Sections were blocked with 1% bovine serum albumin (BSA) for 1.5 h at room temperature and incubated overnight at 4 °C with primary antibody. Immunostaining was performed using the VECTASTAIN Elite ABC IgG kit and NovaRED substrate (Vector Laboratories) according to the manufacturer’s instructions. Images were acquired at ×4 or ×10 magnification using a Nikon Eclipse E200 microscope and Motic Images Plus 3.0 software.

For immunofluorescence, paraffin-embedded mouse, human and porcine wound sections were deparaffinized and subjected to antigen retrieval as previously described(*22*). Sections were blocked for 1 h at room temperature with 5% donkey serum and 1% BSA in PBS containing 0.1% Triton X-100 and incubated overnight at 4 °C in a humidified chamber with primary antibodies. Primary antibodies, dilutions and species-specific applications are listed in **table S4**. Species-appropriate secondary antibodies were used at 1:500 dilution and included Alexa Fluor 488 donkey anti-mouse IgG (Abcam, ab150109), Alexa Fluor 647 donkey anti-rat IgG (Abcam, ab150155), Alexa Fluor 594 donkey anti-rabbit IgG (Abcam, ab150064), Alexa Fluor 488 donkey anti-goat IgG (Abcam, ab150133) and Alexa Fluor 647 donkey anti-goat IgG (Abcam, ab150131). Nuclei were counterstained with DAPI (Invitrogen). Sections were treated with TrueVIEW Autofluorescence Quenching Kit (Vector Laboratories) for 5 min and mounted using VECTASHIELD antifade mounting medium (Vector Laboratories). Images were acquired from multiple fields within the wound area at ×10 or ×20 magnification using a Zeiss Axio Imager A2 or a Zeiss Celldiscoverer 7 operated in widefield mode, with ZEN Blue software. Positively stained single- or double-positive cells were quantified using ImageJ (version 2.1.0) and normalized to the analyzed tissue area.

#### Single-cell preparation and neutrophil fractionation from mouse wounds

Wound tissues were collected from *db/db* mice treated with low-dose Met-gel or BLK-gel and placed in sterile Dulbecco’s phosphate-buffered saline (DPBS; Thermo Fisher Scientific) on ice until processing. Tissues were processed within 3 h of collection. Wound specimens were finely minced using a sterile no. 10 scalpel and digested for 30 min at 37 °C in complete DMEM containing 1.5 mg ml⁻¹ collagenase P (Roche), 1.5 mg ml⁻¹ Dispase II (Gibco) and 1 mg ml⁻¹ DNase I (STEMCELL Technologies). Enzymatic digestion was terminated by addition of complete DMEM containing 10% fetal bovine serum and 1% penicillin-streptomycin. Cell suspensions were passed through 40 μm cell strainers and centrifuged at 500*g* for 10 min at 4 °C. For bulk RNA-sequencing experiments, single-cell suspensions prepared from wounds collected on days 1 and 2 were subjected to magnetic neutrophil enrichment using the Neutrophil Isolation Kit, mouse (Miltenyi Biotec, 130-097-658), according to the manufacturer’s instructions. Non-neutrophil cells were magnetically labelled and retained during separation, whereas unlabeled neutrophils were collected in the flow-through. The resulting neutrophil-enriched and non-neutrophil fractions were collected separately and processed for RNA extraction and bulk RNA sequencing.

#### Flow cytometry

Single-cell suspensions were prepared from day-3 wounds of db/db mice treated with low-dose Met-gel or BLK-gel using the tissue dissociation procedure described above. Cell concentration and viability were assessed using a Cellometer K2 (Nexcelom Bioscience). Dead cells were excluded using a Fixable viability dye (Biolegend, 423107). Single-stained compensation beads (ArC^TM^ Amine Reactive Beads, Invitrogen A10346) were used to establish fluorescence compensation, and unstained cells were used to assist with gating. Cells were blocked for 15 min on ice in FACS buffer containing 5% TruStain FcX^TM^ PLUS (Biolegend, 156604) and 5% True-Stain Monocyte Blocker^TM^ (BioLegend, 426102), followed by surface staining for 30 min on ice with the antibody panels detailed in **table S5**. MR1-5-OP-RU and CD1d-PBS57 tetramers obtained from the NIH Tetramer Core were used to identify MAIT and iNKT cells, respectively. Cells were washed twice and fixed and permeabilized using the Foxp3/Transcription Factor Staining Buffer Set (eBioscience, 00-5521-00) for 45 min on ice. Intracellular staining for CHRM1, IL-17A, FOXP3 and RORγt was performed for 30 min at room temperature. Cells were washed twice with permeabilization buffer (BioLegend, 421002), stored at 4 °C protected from light and acquired the following day on a CytoFLEX LX flow cytometer (Beckman Coulter) at the BIDMC Flow Cytometry Core.

Flow cytometry data were analyzed using CytExpert software version 2.6 (Beckman Coulter). Complete gating strategies for the global immune and innate lymphocyte panels are provided in **fig. S5**. RORγt⁺ MAIT cells were defined as RORγt⁺TCRβ⁺TCRγδ⁻MR1-5-OP-RU tetramer⁺ cells, whereas RORγt⁺ iNKT cells were defined as RORγt⁺TCRβ⁺TCRγδ⁻CD1d-PBS57 tetramer⁺ cells.

#### RNA extraction, cDNA synthesis and quantitative RT-PCR

Total RNA was extracted from isolated cells and wound tissues using the RNeasy Mini Kit (Qiagen, 74014) according to the manufacturer’s instructions. For cellular samples, RNA was isolated from approximately 5x10^5^ cells. For tissue samples, approximately 50 mg of mouse, human or porcine wound tissue was homogenized in QIAzol Lysis Reagent (Qiagen, 79356) using a TissueLyser LT (Qiagen, 85600) before RNA purification with the RNeasy Mini Kit. RNA concentration and purity were assessed using a NanoDrop 2000 spectrophotometer (Thermo Fisher Scientific). cDNA was synthesized from 1 μg total RNA using the RT² First Strand Kit (Qiagen, 330404) according to the manufacturer’s instructions. Quantitative RT-PCR was performed using 10 ng cDNA per reaction with QuantiTect or RT² primer assays (Qiagen) and either the QuantiTect SYBR Green PCR Kit or RT² SYBR Green qPCR Mastermix (Qiagen) on a Stratagene Mx3005P real-time PCR system with MxPro software (Agilent Technologies). Cycling conditions followed the manufacturer’s protocols. Relative gene expression was calculated using the 2^−ΔΔCt^ method. Expression values were normalized to species-specific reference genes: *B2m* for mouse samples, *GAPDH* for human samples and *RPL4* for porcine samples. Primer assay information for mouse, human and porcine targets is provided in **table S6.**

#### Transcriptome sequencing and analysis

Total RNA was isolated from neutrophil-enriched and non-neutrophil fractions obtained from day-1 and day-2 wounds of *db/db* mice treated with BLK-gel or low-dose Met-gel using the RNeasy Plus Universal Mini Kit (Qiagen). RNA quality assessment, library preparation and sequencing were performed by GENEWIZ (South Plainfield, NJ, USA). RNA concentration was determined using a Qubit 4.0 fluorometer (Thermo Fisher Scientific), and RNA integrity was assessed using an Agilent TapeStation. Poly(A)⁺ RNA was enriched using the NEBNext Poly(A) mRNA Magnetic Isolation Module, and sequencing libraries were generated using the NEBNext Ultra II RNA Library Prep Kit for Illumina (New England Biolabs). Library quality and concentration were assessed using an Agilent TapeStation, Qubit fluorometry and quantitative PCR. Libraries were sequenced on an Illumina HiSeq platform using 150-bp paired-end reads, generating approximately 20 million reads per sample. Raw base-call files were converted to FASTQ format and demultiplexed using bcl2fastq (v2.20). Sequencing reads were trimmed to remove adapter sequences and low-quality bases using Trimmomatic (v0.36)(*67*) and aligned to the GRCm38 mouse reference genome using STAR (v2.5.2b)(*68*). Gene-level counts were generated using featureCounts from the Subread package (v1.5.2)(*69*), with uniquely mapped reads overlapping annotated exons included in the count matrix. Differential expression analysis was performed using DESeq2(*70*). *P* values were calculated using the Wald test and adjusted for multiple comparisons using the Benjamini-Hochberg method. Genes with an adjusted *P* < 0.05 and an absolute log₂ fold change >1 were considered differentially expressed. Principal component analysis was performed using the plotPCA function in DESeq2. Functional enrichment analysis was performed using Metascape(*71*). Significantly upregulated and downregulated genes were submitted as separate gene lists, and enriched biological processes and pathways were identified using the integrated ontology resources implemented in Metascape.

To assess treatment-dependent changes in early transcriptional trajectories, differential expression between day 2 and day 1 was calculated separately for BLK-gel- and Met-gel-treated wounds. Analyses were restricted to significantly differentially expressed genes, using the criteria defined above, and genes were ranked by the magnitude of their log₂ fold change. For each gene, the day 2 versus day 1 log₂ fold change under BLK-gel treatment was compared with the corresponding temporal log₂ fold change under Met-gel treatment. Genes exhibiting temporal changes in opposite directions between treatments were classified as having reversed trajectories. Representative genes showing the largest opposing changes were highlighted in trajectory plots and visualized as heat maps. Selected significant genes were additionally grouped by functional annotation to illustrate treatment-associated changes in inflammatory signaling, chemotaxis, granulocyte activation and tissue remodeling processes. Day-1 treatment-responsive genes were visualized separately using DESeq2-derived log₂ fold changes for Met-gel versus BLK-gel treatment. Analyses were performed separately for the neutrophil-enriched and non-neutrophil fractions. Downstream data processing and visualization were performed in Python using pandas, NumPy and Matplotlib.

#### Xenium spatial transcriptomic profiling

Formalin-fixed, paraffin-embedded (FFPE) wound tissues from Met-gel- and BLK-gel-treated *db/db* mice collected on day 3 were processed for targeted spatial transcriptomic profiling by the BIDMC Spatial Technologies Unit. Sample preparation, tissue processing, probe hybridization, amplification, imaging and transcript decoding were performed according to the BIDMC TMC/STU Xenium methodological workflow(*72*). Eight wound samples (*n* = 4 per treatment group) were analyzed using a fully custom-designed 471-gene Xenium panel selected to characterize immune, stromal, epithelial and vascular cell populations and transcriptional programs associated with cholinergic signaling, IL-17 signaling and wound repair. The complete 471-gene panel is provided in **table S2**.

#### Spatial Transcriptomic Data Processing and Analysis

##### Cell segmentation, clustering and annotation

Cell segmentation was performed across eight Xenium samples using the transcript-based segmentation method Proseg(*73*). Segmented transcript counts and cell-level metadata were imported into Seurat (v5.4.0)(*74*). Gene expression data were normalized and variance stabilized using SCTransform with v2 regularization. Principal component analysis (PCA) was performed on variable features, and sample-associated batch effects were corrected in PCA space using Harmony (v1.2.3)(*75*), with sample identity specified as the batch variable. A shared nearest-neighbor graph was constructed using the first 15 Harmony-corrected principal components, and graph-based clustering was performed using the Louvain algorithm at a resolution of 0.3. Cluster-enriched marker genes were identified by differential expression analysis and used to manually annotate clusters into eleven major cell types. Low-dimensional representations and cell-type distributions were visualized using UMAP.

##### Anatomical region annotation

Anatomical regions were manually delineated for each sample using the tissue morphology images in Xenium Explorer (v4.0; 10x Genomics). Regions were classified as non-wound tissue, wound center, wound edge, scab or migrating epidermis. Region annotations were subsequently incorporated into downstream spatial analyses.

##### Spatial neighborhood analysis

Spatial neighborhood analyses were performed separately for Met-gel- and BLK-gel-treated samples and within each manually annotated anatomical region. For each sample and region, a spatial neighbor graph was constructed from cell-centroid coordinates using a fixed interaction radius of 25 μm, excluding each cell itself. For each cell, neighborhood composition was calculated as the number of neighboring cells of each annotated cell type and normalized to the total number of cells within its neighborhood. Cells without neighbors within 25 μm were excluded. Cellular neighborhoods were identified by mini-batch *k*-means clustering of the resulting per-cell neighborhood-composition vectors (*k* = 15). For each cellular neighborhood, cell-type enrichment was calculated as the log₂ ratio of its meancell-type composition to the corresponding cell-type abundance within that anatomical region.

##### NK-cell microenvironment analysis

To characterize the cellular microenvironment surrounding NK cells, neighborhood-composition vectors corresponding to annotated NK cells were extracted from the spatial neighbor graphs. Neighborhood compositions were first averaged within each sample and subsequently across samples within each anatomical region, thereby preventing samples containing larger numbers of NK cells from disproportionately contributing to the group-level estimate. NK-neighborhood enrichment for each cell type was calculated relative to its overall abundance within the corresponding anatomical region. Enrichment values greater than 1 indicated cell types occurring more frequently within 25 μm of NK cells than expected from their regional abundance.

##### Gene-signature activity analysis

Gene-signature activity was quantified at single-cell resolution using custom gene sets representing CHRM1- and IL-17-associated signaling. The CHRM1-associated signature comprised *Chrm1, Rgs2, Grk2, Plcb1, Plcb3, Plcb4, Plcg1, Plcg2, Itpr1, Plcd1, Prkca, Prkcb, Camk2d, Calm1, Stim1, Orai1, Nfatc1, Nfatc2, Nfatc3, Mapk1, Mapk3, Mapk14, Mapk8, Mapk9, Nr4a1, Nr4a2* and *Fosl1*. The IL-17-associated signature comprised *Il17a, Il17f, Il17ra, Il17rc, Ccl20, Cxcl15, Cxcl10, Nfkb1, Nfkb2, Rela, Il6, Il1a* and *Mmp3*. For each cell, genes were ranked according to expression, and the resulting ranks were centered and scaled to generate empirical z-ranks. Positive and negative ranks were processed separately, and technical effects associated with the number of detected genes and total UMI counts were removed by linear regression. Signature scores were calculated from the resulting rank-transformed expression values for genes belonging to each custom signature. The resulting signature-score matrix was further corrected for sample-associated batch effects and residual technical variation associated with the number of detected genes and log-transformed total UMI counts using removeBatchEffect from the limma package(*58*).

##### Differential pathway activity between conditions

Differences in pathway activity between treatment groups (Drug versus Control) were assessed per cell type using Limma. For each cell type, log fold-changes were extracted for every pathway.

##### NK cell ligand-receptor inflow analysis

We performed NK-receptor ligand receptor signaling using inflow score implemented in LIANA+(*76*). Analyses were performed for each region (non-wound, wound center, wound edge, scab, and migrating epidermis) per sample. The LIANA consensus ligand-receptor resource was converted to mouse orthologs using decoupler(*77*), retaining mappings supported by at least three evidence and allowing up to two one-to-many ortholog mappings. Only ligand-receptor pairs represented in the Xenium gene panel were evaluated.

LIANA inflow scores were computed for each section independently. Spatial neighborhoods were constructed using a bandwidth of 200, and samples with fewer than 50 cells were excluded. NK-cell communication was evaluated in two directions. For incoming signaling, inflow scores were averaged across NK cells to quantify ligand-receptor interactions originating from other cell types and received by NK cells. For outgoing signaling, interactions in which NK cells served as the ligand-producing source were summarized across cells of each receiving cell type. Per Sample inflow scores were computed separately for each region, interacting partner cell type, and ligand-receptor pair using two-sided Mann-Whitney U tests to assess differential communication between control and drug conditions. Effect sizes were summarized as the log2 fold change of mean inflow scores, and multiple-testing correction was performed independently within each spatial region using the Benjamini-Hochberg procedure.

#### Western blotting

Wound tissues collected from mice on day 5 were weighed and homogenized in T-PER Tissue Protein Extraction Reagent (Thermo Fisher Scientific) supplemented with protease and phosphatase inhibitor cocktails (Thermo Fisher Scientific, 78430 and 78420) using a TissueLyser LT (Qiagen) at 50 Hz. Homogenates were incubated on ice for 30 min with intermittent vortexing and centrifuged at 14,000*g* for 30 min at 4 °C. Supernatants were collected, and total protein concentration was determined using the Pierce BCA Protein Assay Kit (Thermo Fisher Scientific). Protein lysates (20 μg per sample) were mixed with 6× Laemmli sample buffer and denatured at 95 °C for 5 min. Proteins were separated on 12% SDS-PAGE gels (Bio-Rad) at 100 V for 45 min and transferred to membranes at 60 V for 3 h at 4 °C. Membranes were blocked in 5% bovine serum albumin (BSA) in Tris-buffered saline containing Tween-20 (TBST) and incubated overnight at 4 °C with primary antibodies against Vinculin, MMP-9, TIMP-1 and NGAL (Santa Cruz Biotechnology; 1:1,000). Membranes were washed with TBST and incubated with the appropriate secondary antibodies (1:10,000) for 1 h at room temperature. Protein bands were detected using a chemiluminescent substrate (Bio-Rad) and imaged using a ChemiDoc Touch Imaging System (Bio-Rad). Band intensities were quantified by densitometry using ImageJ (v2.1.0), and expression of each target protein was normalized to Vinculin.

#### Multiplex cytokine and chemokine analysis

Mouse wound tissues collected on day 3 were homogenized as described above, and tissue lysates were adjusted to a total protein concentration of 3 mg ml⁻¹. Samples were analyzed by Eve Technologies (Calgary, Canada) using the Mouse Cytokine/Chemokine 44-Plex Discovery Assay (MD44). Wound exudates collected on day 4 from treated wounds in diabetic Yucatan miniature swine by centrifugation of the treatment hydrogel dressings and were analyzed by Eve Technologies using the Porcine Cytokine/Chemokine 13-Plex Discovery Assay (PD13).

#### Acetylcholine and acetylcholinesterase measurements

Wound tissue lysates prepared as described above were used to quantify acetylcholine concentrations and AChE activity. Acetylcholine concentrations were measured using the Acetylcholine Assay Kit (Sigma-Aldrich, MAK435) according to the manufacturer’s instructions, with absorbance measured at 570 nm. AChE activity was measured using the Acetylcholinesterase Inhibitor Screening Kit (Sigma-Aldrich, MAK324) according to the manufacturer’s instructions, with absorbance measured at 412 nm. Acetylcholine concentrations and AChE activity were normalized to total protein content determined using the Pierce BCA Protein Assay Kit (Thermo Fisher Scientific).

#### *Ex vivo* human skin wound healing model

De-identified human skin specimens were obtained from Genoskin Inc (Salem, MA) for *ex vivo* wound healing studies. Donor tissues were collected by the provider in accordance with applicable ethical and regulatory requirements and were screened for infectious agents, including hepatitis B virus, hepatitis C virus, HIV-1 and HIV-2. Donor characteristics for all human skin experiments are provided in **table S7**. Full thickness human skin was prepared as 1-cm-diameter circular explants, and a 2 mm wound was created at the center of each explant. Immediately after wounding, metrifonate solution (Met-sol;10ul at 1 mg ml⁻¹) or PBS vehicle control was applied directly to the wound area. Explants were maintained in culture medium provided by Genoskin according to the manufacturer’s instructions and imaged daily from day 0 to day 4. Wound area was quantified from serial macroscopic images using ImageJ (v2.1.0) by manually delineating the wound boundary defined by the leading edge of migrating keratinocytes. Wound area at each time point was expressed as a percentage of the corresponding day 0 wound area. Conditioned culture media were collected on days 2 and 4 for cytokine analysis. On day 4, skin tissues were collected for histological and immunofluorescence analyses.

#### ELISA assay

IL-17A concentrations in conditioned media from *ex vivo* human skin wounds and wound exudates from diabetic Yucatan miniature swine were quantified using species-specific ELISA kits according to the manufacturers’ instructions (human IL-17A High Sensitivity ELISA, Invitrogen, BMS2017-2HS; porcine IL-17A ELISA, Invitrogen, ESIL17A). Conditioned media from human skin explants were collected on days 2 and 4 after wounding and concentrated using 10-kDa molecular-weight-cut-off centrifugal filters (Amicon) in the presence of protease inhibitor cocktail (Thermo Fisher Scientific). Porcine wound exudates were collected on day 4 after wounding. Human conditioned-media concentrates and porcine wound exudates were assayed without further dilution. Samples and recombinant protein standards were added to antibody-coated 96-well plates, and ELISAs were performed according to the respective manufacturer’s protocol. Absorbance was measured at 450 nm using a microplate reader, and IL-17A concentrations were calculated from standard curves generated using serial dilutions of the corresponding recombinant protein standards. IL-17A concentrations were normalized to total protein content determined using the Pierce BCA Protein Assay Kit (Thermo Fisher Scientific).

#### Statistical analysis

Statistical analysis was performed using GraphPad Prism (v10) and Minitab (v22.3.1). For repeated-measures data involving multiple experimental groups, a mixed-effects model was applied with day and treatment as fixed effects. Post hoc comparisons were performed using Fisher’s least significant difference (LSD) multiple-comparison test to identify significant differences between groups at individual time points. For comparisons between two groups, two-sided unpaired *t*-tests were used. Data are presented as mean ± standard deviation (SD), as indicated in the figure legends. *P* values < 0.05 were considered statistically significant, and all tests were performed using the default settings of the software unless otherwise specified.

**Fig. S1.**
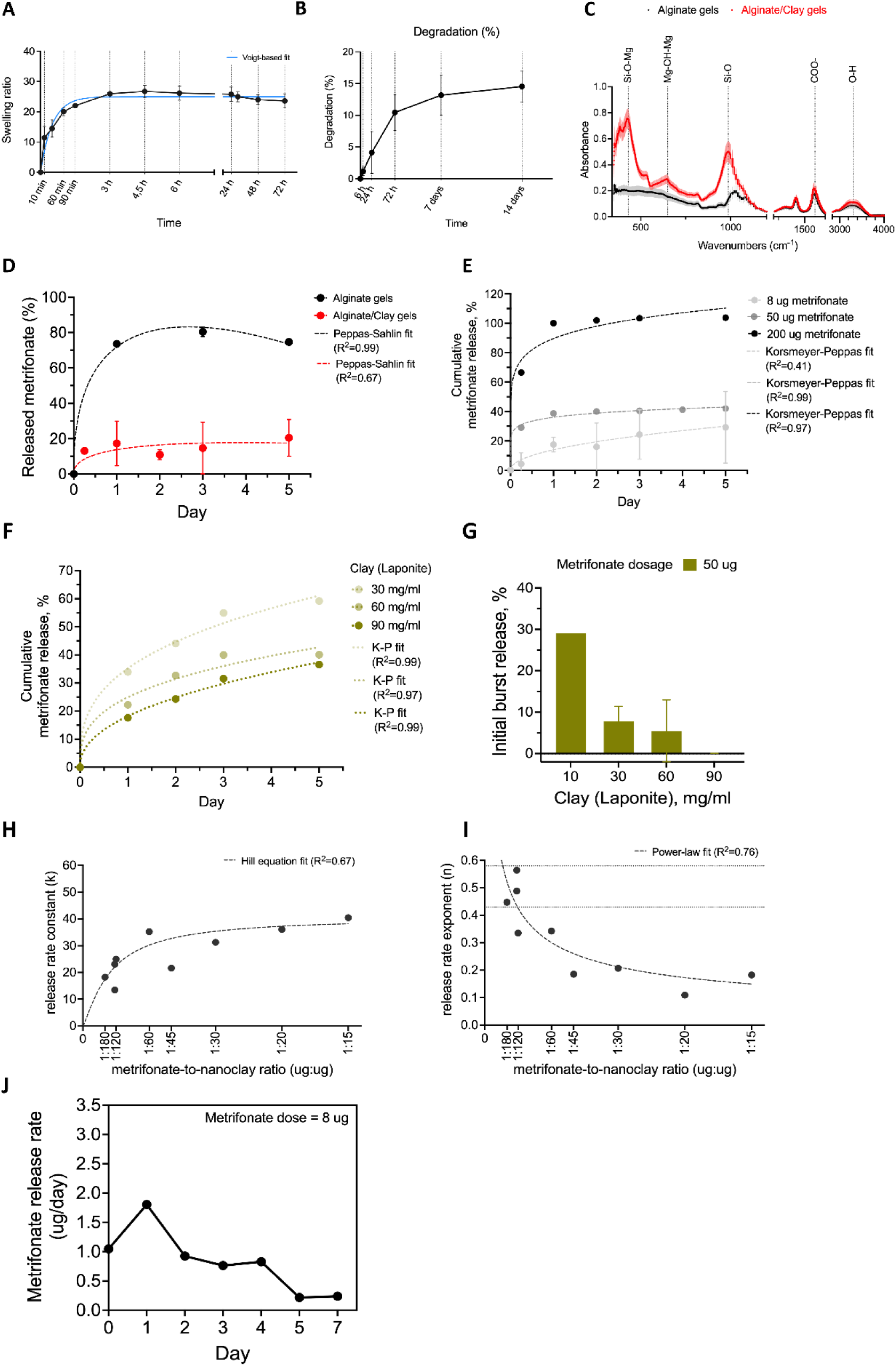
Characterization of metrifonate-loaded alginate/clay hydrogel (Met-gel). **A,** Swelling ratio of Met-gels as a function of incubation time in PBS. The data points are fitted to the Voigt-based model, which is indicated by a solid blue line. **B,** Percentage degradation of Met-gels over time in PBS, as quantified by mass loss compared to day 0. Data represent mean ± SD for n=3 hydrogels per condition. **C,** Fourier Transform Infrared Spectroscopy (FTIR) spectrum of alginate gels (black curve) and alginate/clay gels (red curve), showing characteristic alginate bands corresponding to COO^-^ and O-H stretching at approximately 1600 and 3300 cm^-1^, respectively, and characteristic nanoclay bands corresponding to Si-O-Mg, Mg-OH-Mg, and Si-O vibrations at approximately 450, 650, and 987 cm^-1^ respectively. **D,** Metrifonate release from alginate hydrogels with (red line) or without clay nanoparticles (black line). Alginate/Clay gels contained 8 µg metrifonate and 10 mg/mL laponite. **E,** Effect of metrifonate loading (8, 50, or 200 µg) on cumulative release from alginate/clay hydrogels; 10 mg/mL of laponite was encapsulated in all gels. **F,** Effect of clay nanoparticles concentration (30, 60, or 90 mg/mL) on cumulative release from alginate/clay hydrogels loaded with 50 µg metrifonate. **G,** Initial burst release of metrifonate on day 1 as a function of clay nanoparticles concentration for gels loaded with 50 µg metrifonate. The data points were fitted to the Peppas-Sahlin model **(D)** and the Korsmeyer-Peppas (K-P) model **(E, F)**, with the resulting curves indicated by dashed lines. Data represent mean ± SD for n=2 or 3 hydrogels per condition. **H,** Correlation with the release rate constant (). The data points were fitted to a Hill equation (dashed line). The positive, plateauing relationship indicates that as the amount of loaded metrifonate increases relative to the nanoclay, the matrix binding capacity approaches saturation, leading to higher release rates, and potentially causing a burst release. **I,** Correlation with the release rate exponent (). The data points were fitted to a power-law model (dashed line), demonstrating a negative relationship. As the amount of loaded metrifonate decreases relative to the nanoclay, the n value falls into the 0.43-0.58 range (highlighted by horizontal dotted lines), indicating that the drug release mechanism becomes dominated by Fickian diffusion. **J,** Daily release of Low Dose (LD) metrifonate/clay hydrogel by weight over 7 days.

**Fig. S2.**
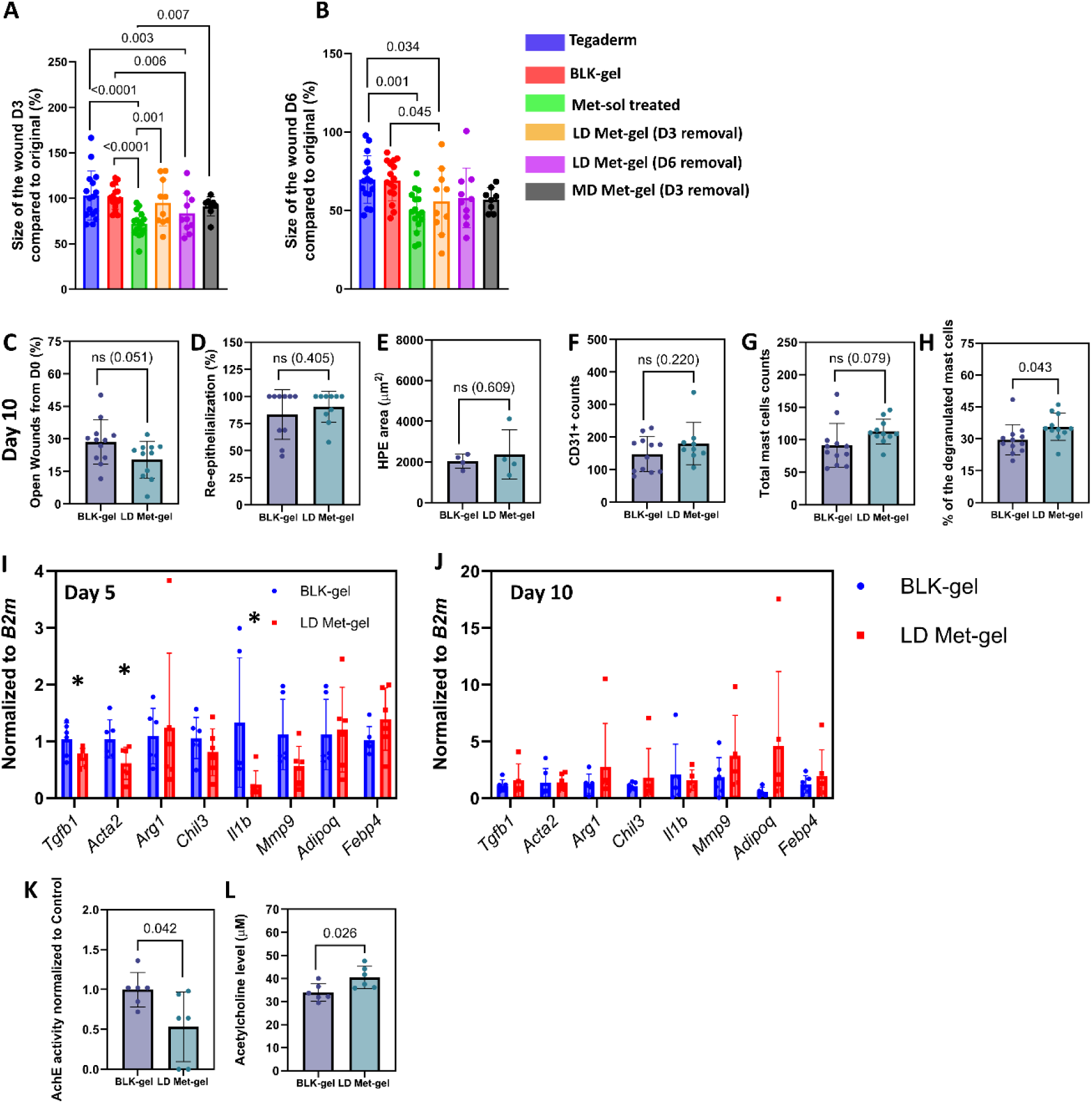
Evaluation of multiple healing metrics in wounds treated with Met-gel. **A-B,** Quantification of the wound closure on Day 3 (**A**), and Day 6 (**B**). Data are presented as mean ± sd, (*n*=10-16). *P* values were determined using mixed-effects model with Days and Treatment as fixed effects, followed by Fisher’s LSD multiple-comparison test. **C,** Quantification of the wound closure on Day 10. **D,** Quantification of re-epithelialization on Day 10 expressed as %. **E,** The hyperproliferative epidermal (HPE) area on Day 10. **F,** Quantification of CD31+ vessels on Day 10. **G,** Quantification of total mast cells on Day 5. **H,** the proportion of degranulated mast cells. Data in **C-H** presented as mean ± s.d. (*n* = 10-12 wounds). *P* values were determined using two-sided unpaired *t*-test. ns, not-significant (*P* ≥ 0.05) **I-J,** RT-qPCR gene expression of mouse wounds on Day 5 (**I**) and Day 10 (**J**). Data are presented as mean ± s.d. (*n* = 6 wounds). *P* values were determined using two-sided unpaired *t*-test. *, *P* ≤ 0.05. **K**, Acetylcholinesterase activity in mouse wounds on D5. **L**, Acetylcholine levels in mouse wounds on D5. Data in **k** and **l** are presented as mean ± s.d. (*n* = 6 wounds). *P* values were determined using two-sided unpaired *t*-test.

**Fig. S3.**
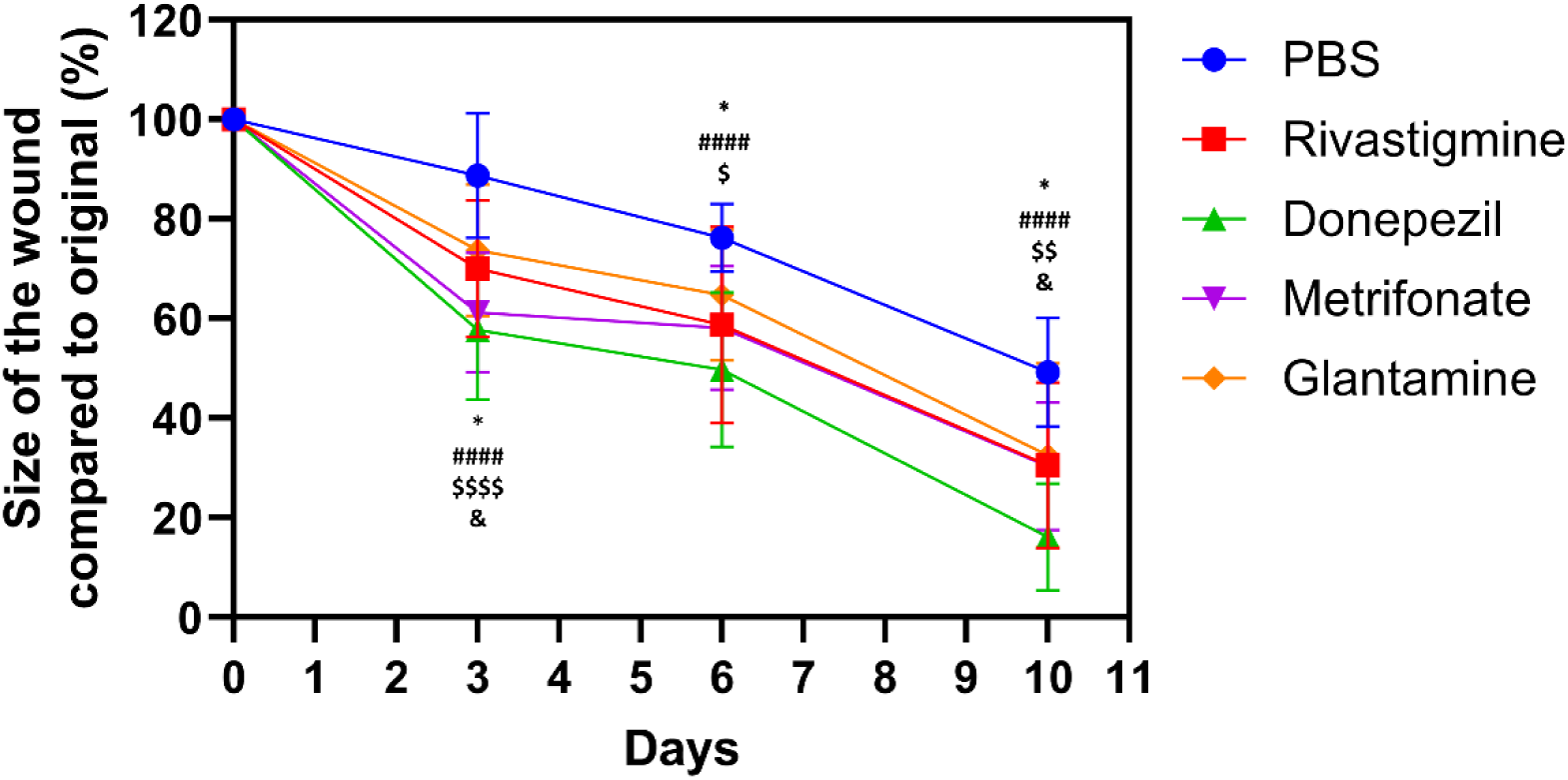
Evaluation of clinically used AChE inhibitors, Rivastigmine, Donepezil, Metrifonate, and Galantamine in *db/db* mouse wound healing. Data are mean ± s.d. (*n*=10 wounds) *: PBS vs Rivastigmine, * (*P*<0.05); #: PBS vs Donepezil, #### (*P*<0.0001); $: PBS vs Metrifonate, $ (*P*<0.05), $$ (*P*<0.01), $$$$ (*P*<0.0001); &: PBS vs Glantamine, & (*P*<0.05). *P* values were determined using mixed-effects model with Days and Treatment as fixed effects, followed by Fisher’s LSD multiple-comparison test.

**Fig. S4.**
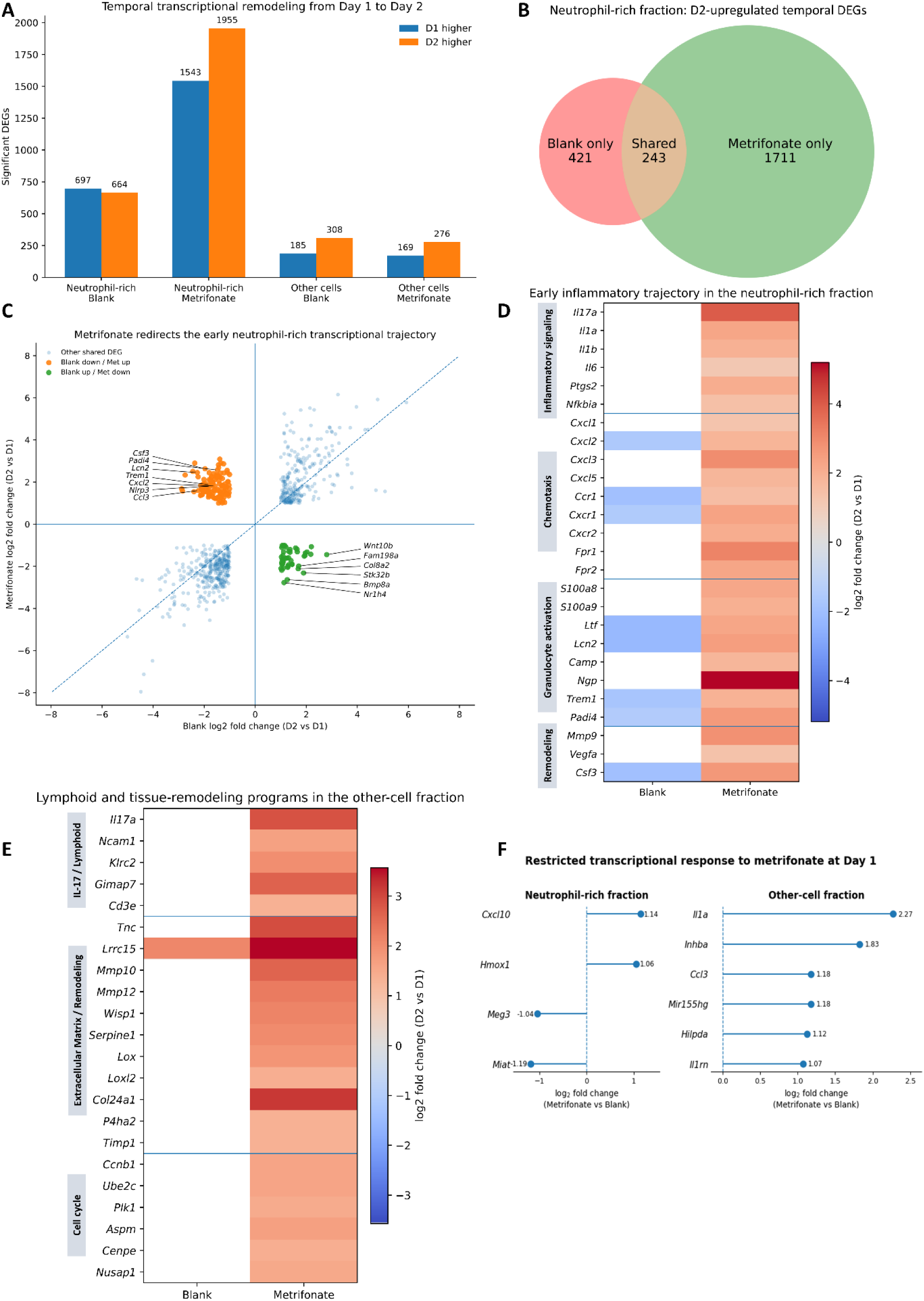
Cholinergic modulation redirects the early transcriptional trajectory of diabetic wounds. **A,** Number of significantly differentially expressed genes (DEGs) between day 1 and day 2 after wounding in neutrophil-rich and other-cell fractions isolated from blank gel- and metrifonate gel-treated diabetic wounds. Genes with positive log2 fold change were more highly expressed at day 2, whereas genes with negative log2 fold change were more highly expressed at day 1. **B,** Venn diagram showing the overlap of genes significantly upregulated at day 2 relative to day 1 in neutrophil-rich fractions from blank gel- and metrifonate-treated wounds. **C,** Comparison of day 1-to-day 2 transcriptional changes in neutrophil-rich fractions. Each point represents a gene significantly differentially expressed over time in both treatment groups. Genes that decreased from day 1 to day 2 in blank gel-treated wounds but increased following metrifonate treatment are highlighted, together with genes displaying the reciprocal trajectory. Selected genes associated with inflammatory/myeloid activation are indicated. **D,** Temporal changes in selected genes associated with IL-17/cytokine signaling, chemotaxis, granulocyte activation and repair in neutrophil-rich fractions. **E,** Temporal changes in selected IL-17/lymphoid, extracellular matrix/remodeling and cell-cycle-associated genes in the other-cell fraction. Heatmaps in **D, E** show log2 fold change for day 2 relative to day 1. **F,** Ranked dot plots showing significantly DEGs in the neutrophil-rich and other-cell wound fractions at Day 1 after wounding. Differential expression analyses were performed independently within each treatment and cellular fraction.

**Fig. S5.**
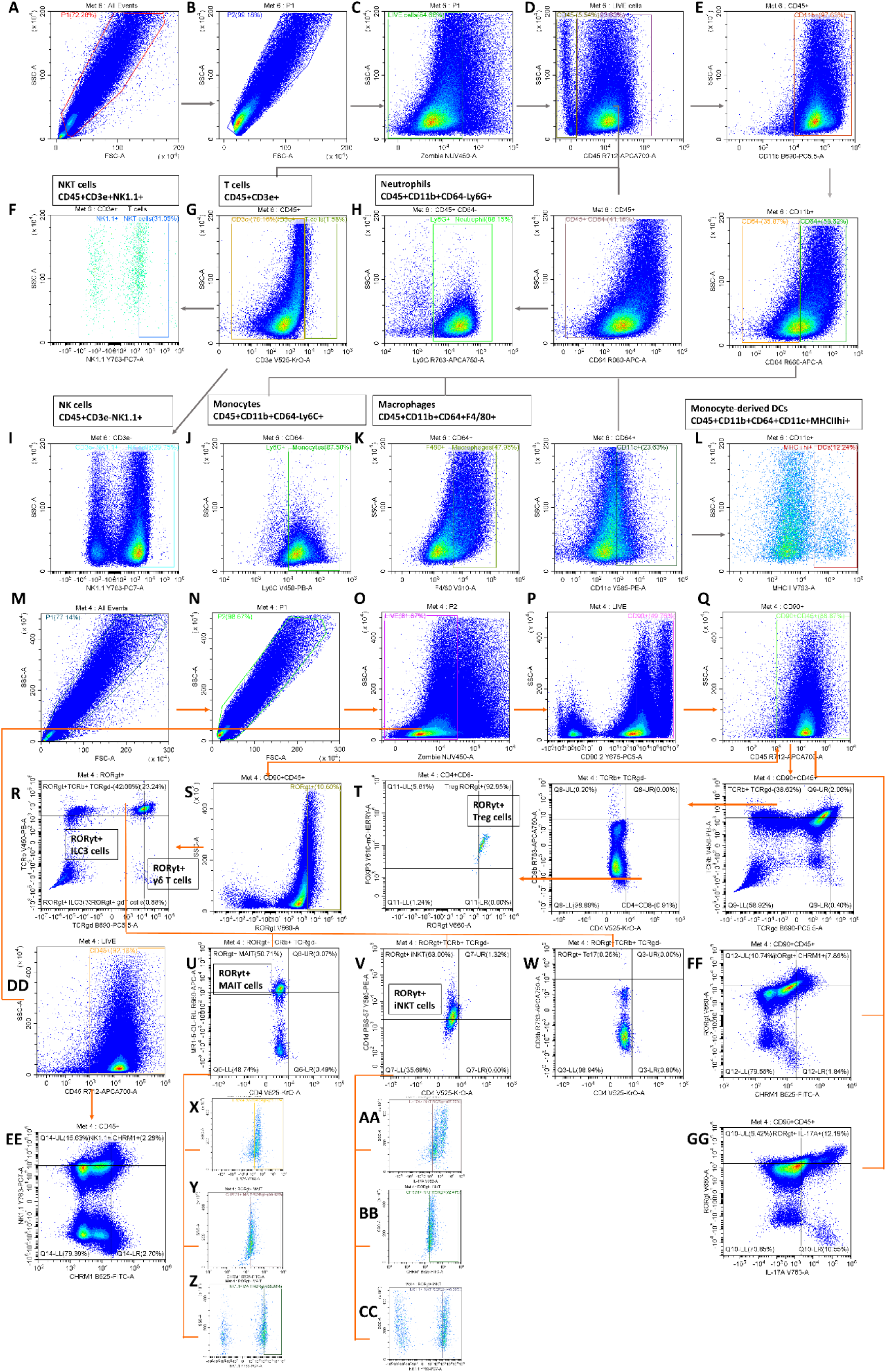
Flow cytometry gating strategy for global immune cell populations and subclassification of RORγt⁺ cells from dissociated wound tissues treated LD Met-gel or Blk-gel. Debris exclusion (A) and doublet exclusion (B) were performed before gating for Live (C), and Live CD45+ population (D). Natural Killer (NK)T cells (CD45+CD3e+NK1.1+) (F), T cells (CD45+CD3e+) (G), Neutrophils (CD45+CD11b+CD64-Ly6G+) (H), and NK cells (CD45+CD3e-NK1.1+) (I) were directly gated from Live CD45+. Monocytes (CD45+CD11b+CD64-Ly6C+) (J), Macrophages (CD45+CD11b+CD64+F4/80+) (K), and Monocyte-derived Dendritic Cells (DCs) (CD45+CD11b+CD64+CD11c+MHCIIhi+) (L) are gated within Live CD45+CD11b+ (E). Debris exclusion (M) and doublet exclusion (N) were performed before gating for Live (O), and Live CD90+CD45+ population (P, Q). CD90+CD45+ RORγt+ cells (S) were then characterized as ILC3 cells (TCRγδ-TCRβ-), γδ T cells (TCRγδ+TCRβ-) (R), Treg cells (TCRγδ-TCRβ+CD8-CD4+FOXP3+) (T). MAIT cells (MR1-5-OP-RU tetramer+CD4-) (U) and iNKT cells (CD1d PBS-57 tetramer+CD4-) (V), and Tc17 cells (CD4-CD8+) (W) were both characterized within RORγt+TCRγδ-TCRβ+ population of cells (R). The Fraction of cells expressing IL-17A+ (X, AA), CHRM1+ (Y, BB), and NK1.1 (Z, CC) are gated for MAIT cells and iNKT cells. The proportion of NK1.1+ CHRM1+ cells (EE) are gated from Live CD45+ (DD). The proportion of RORγt+CHRM1+ cells (FF) and RORγt+IL-17A+ cells (GG) are gated from Live CD90+CD45+ (Q).

**Fig. S6.**
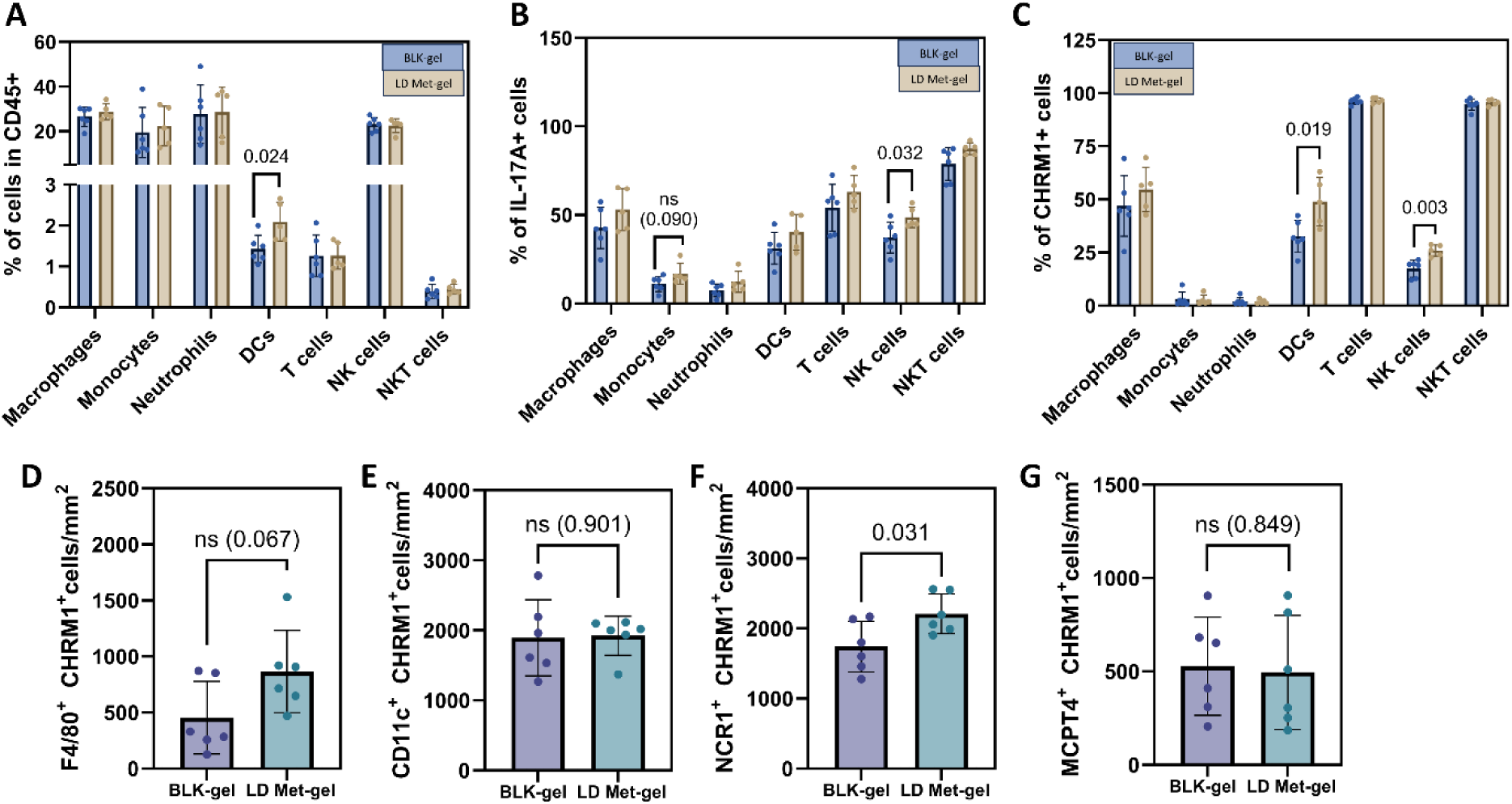
Broad phenotypic analysis of immune cell populations in Day 3 wounds. The proportion of macrophages (CD45+CD11b+CD64+F4/80+), monocytes (CD45+CD11b+CD64-Ly6C+), neutrophils (CD45+CD11b+CD64-Ly6G+), monocyte derived dendritic cells (DCs) (CD45+CD11b+CD64+CD11c+MHCIIhi+), T cells (CD45+CD3e+), natural killer (NK) cells (CD45+CD3e-NK1.1+) and NKT cells (CD45+CD3e+NK1.1+) are gated on Live CD45+(**A**). The Fraction of cells expressing IL-17A (**B**) and CHRM1 (**C**) are quantified from each cell population. **D-G,** Immunofluorescence quantitation of CHRM1 expression in: macrophages - F4/80+cells (**D**), dendritic cells - CD11c+cells (**E**), NK cells - NCR1+cells (**F**), and mast cells - MCPT4+ cells (**G**) per unit area. Data in **A-G** are presented as mean ± s.d. (*n* = 5-6). *P* values were determined using two-sided unpaired *t*-test.

**Fig. S7.**
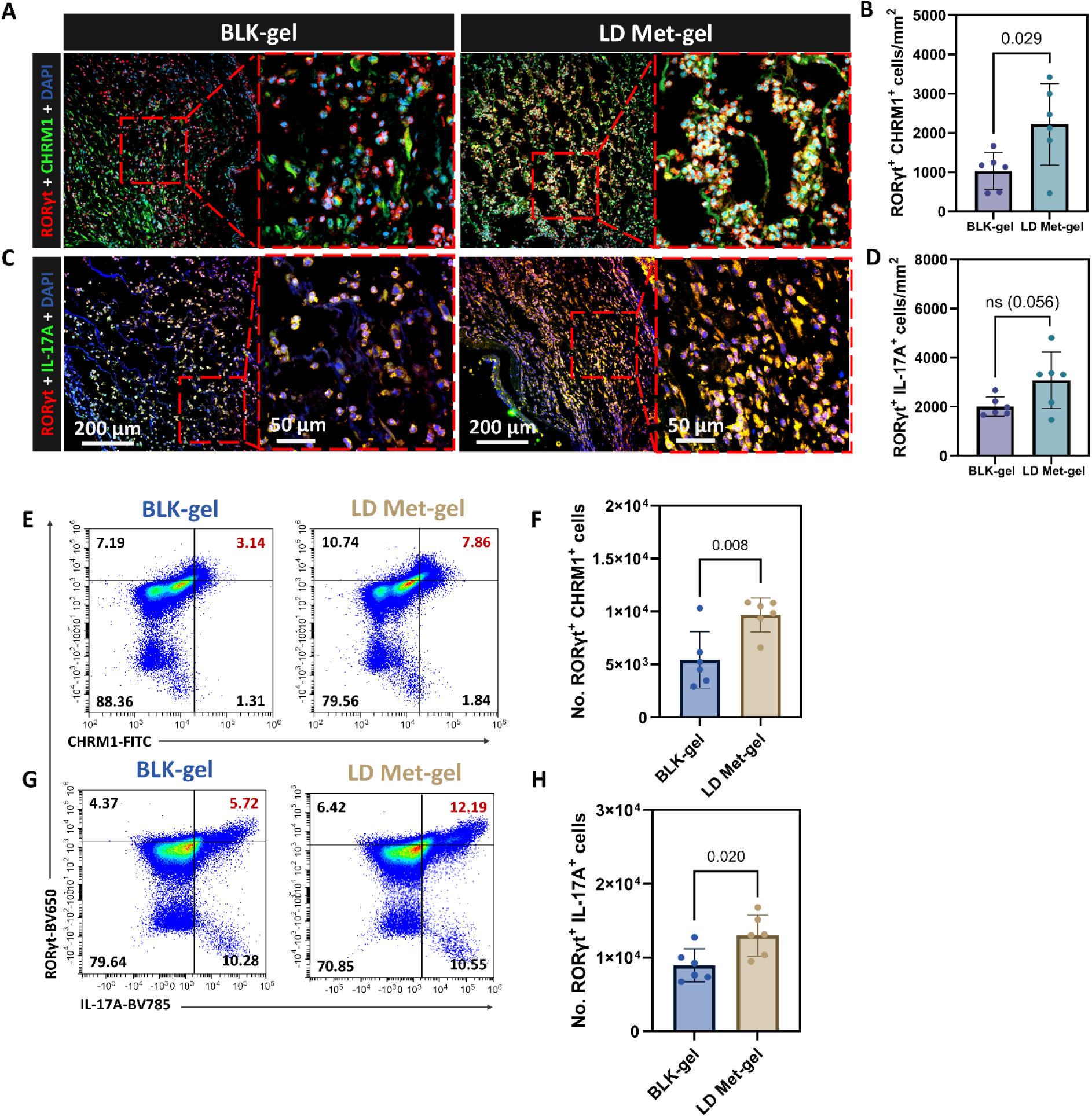
Cholinergic modulation increases RORγt⁺CHRM1⁺ and RORγt⁺IL-17A⁺ cells in diabetic wounds. **A**, Representative immunofluorescence images for RORγt & CHRM1 and **B,** Quantification of RORγt+ CHRM1+ double positive cells per unit area. **C**, Representative immunofluorescence images for RORγt & IL-17A and **D,** Quantification of RORγt+ IL-17A+ double positive cells per unit area. **E,** Representative flow cytometry scatter plots of RORγt and CHRM1 (**E**), and RORγt and IL-17A (**G**), gated on CD90+CD45+ cells. Quantification of RORγt+ CHRM1+ cells gated on live cells (**F**), and RORγt+ IL-17A+ cells gated on live cells (**H**) Data in **B, D, F, H** are presented as mean ± s.d. (*n* = 6). *P* values were determined using two-sided unpaired *t*-test. Histology samples and single-cell suspensions were obtained from *db/db* mice dorsum wounds on day 3.

**Fig. S8.**
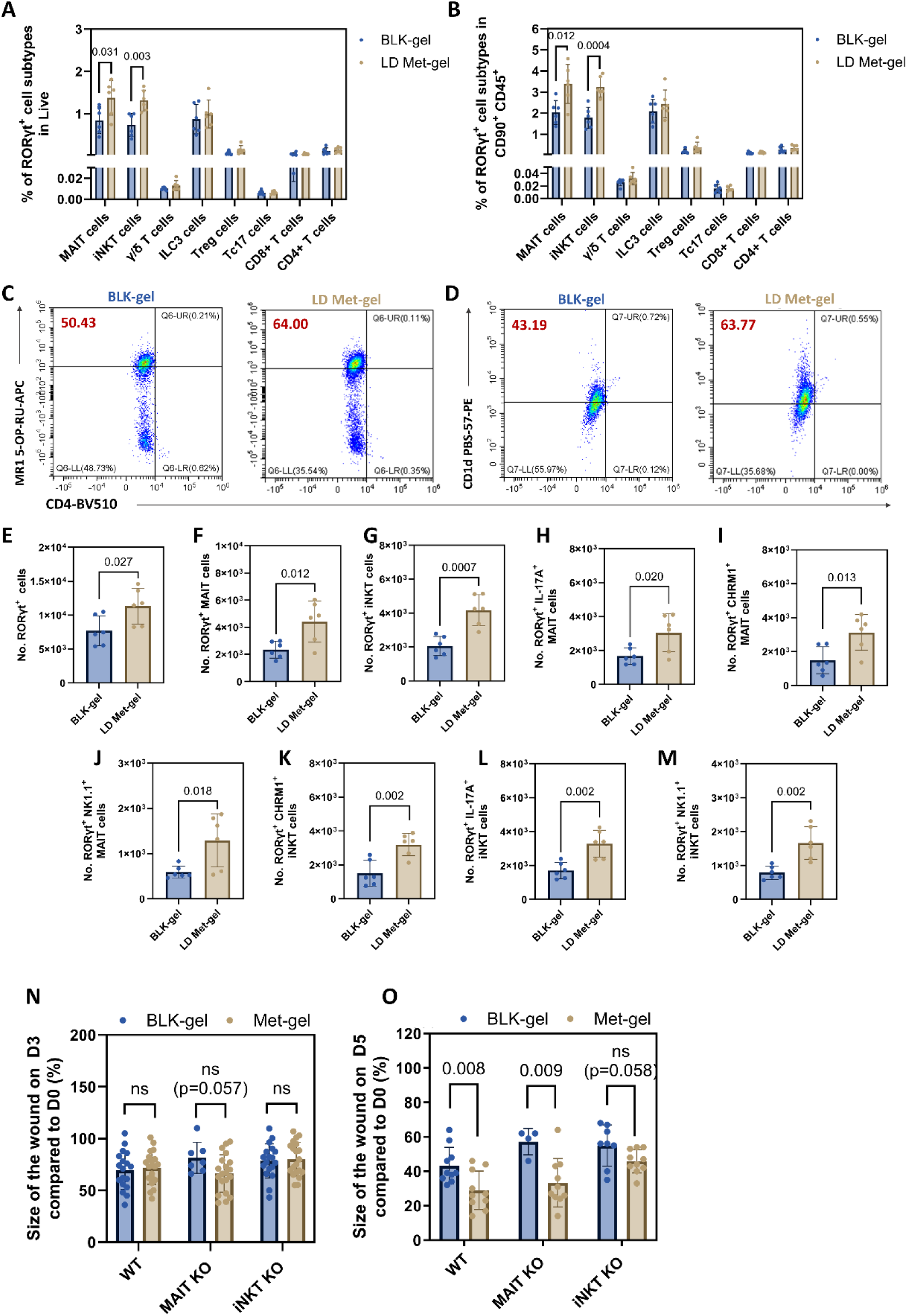
MAIT and iNKT cells exhibit treatment-associated phenotypic changes but are dispensable for enhanced wound repair following cholinergic modulation. The proportion of RORγt⁺ cells subsets are gated on live (A), and Live CD45+CD90+ (B). Representative flow cytometry plots for the population of MAIT cells (MR1-5-OP-RU tetramer+CD4-) (C), and the population of iNKT cells (CD1d PBS-57 tetramer+CD4-) in wounds treated with LD Met-gel or BLK-gel and they are both gated within CD45+CD90+RORγt+ (D). Total number RORγt+ cells (E), RORγt+MAIT cells (F), RORγt+iNKT cells (G) in wounds treated with LD Met-gel or BLK-gel. IL-17A, CHRM1, and NK1.1 expression are quantified on both MAIT cells and iNKT cells (H-M). Data in A-B and E-N are presented as mean ± s.d. (*n* = 6). *P* values were determined using two-sided unpaired *t*-test. Single-cell suspensions were obtained from *db/db* mice dorsum wounds on day 3. N-O, Wound healing in LD Met and BLK-gel treated STZ induced Wild Type (WT - C57BL/6J), MAIT cell Knockout (KO) mice (*Mr*1-/-) and iNKT cell KO mice (*Traj*18-/-) on day 3 **(N)** and day 5 **(O)**. Data are presented in **N** and **O** as mean ± s.d. (*n* = 4-20) were analyzed using two-sided unpaired *t*-test.

**Fig. S9.**
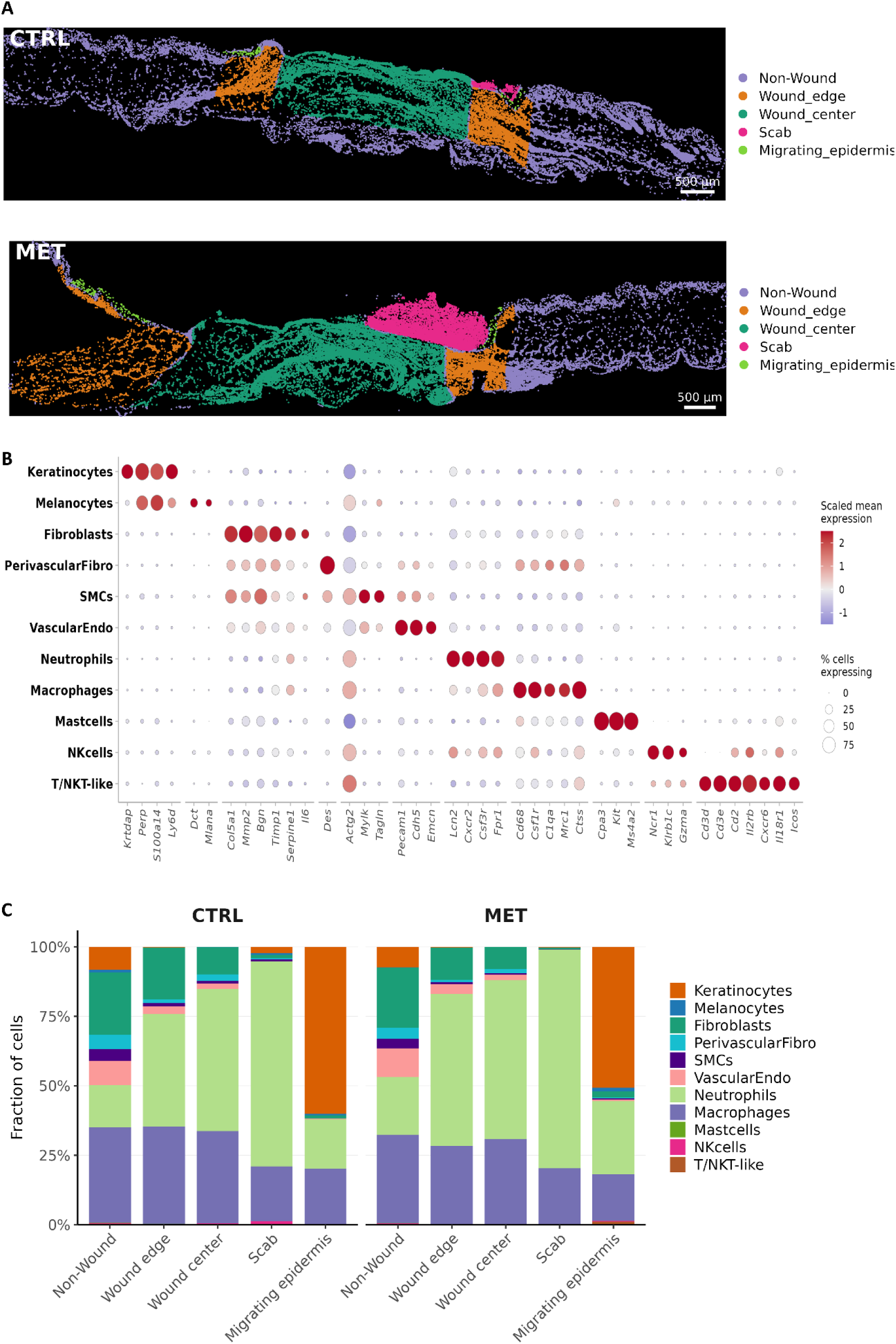
Spatial annotation and cellular composition of *db/db* mouse wounds profiled by Xenium. **A**, Representative spatial maps of day 3 diabetic wounds treated with blank gel (CTRL) or metrifonate gel (MET), showing annotation of the wound regions used for subsequent spatial analyses, including the wound edge (orange), wound center (green), scab (pink), migrating epidermis (bright green) and non-wounded area (purple). **B**, Cell-type annotation of Xenium spatial transcriptomic data. Dot plot showing the expression of representative cell-type marker genes across 11 annotated cell populations, including keratinocytes, melanocytes, fibroblasts, perivascular fibroblasts, smooth muscle cells (SMCs), vascular endothelial cells, neutrophils, macrophages, mast cells, natural killer (NK) cells, and T/NKT-like cells. Dot size represents the percentage of cells expressing each gene, and color indicates scaled mean gene expression within each cell population. **C**, Distribution and relative abundance of transcriptionally defined cell clusters across individual BLK-gel (CTRL)- and Met-gel (MET)-treated wounds. Clusters were assigned to major cell identities based on established lineage and cell-type marker expression.

**Fig. S10.**
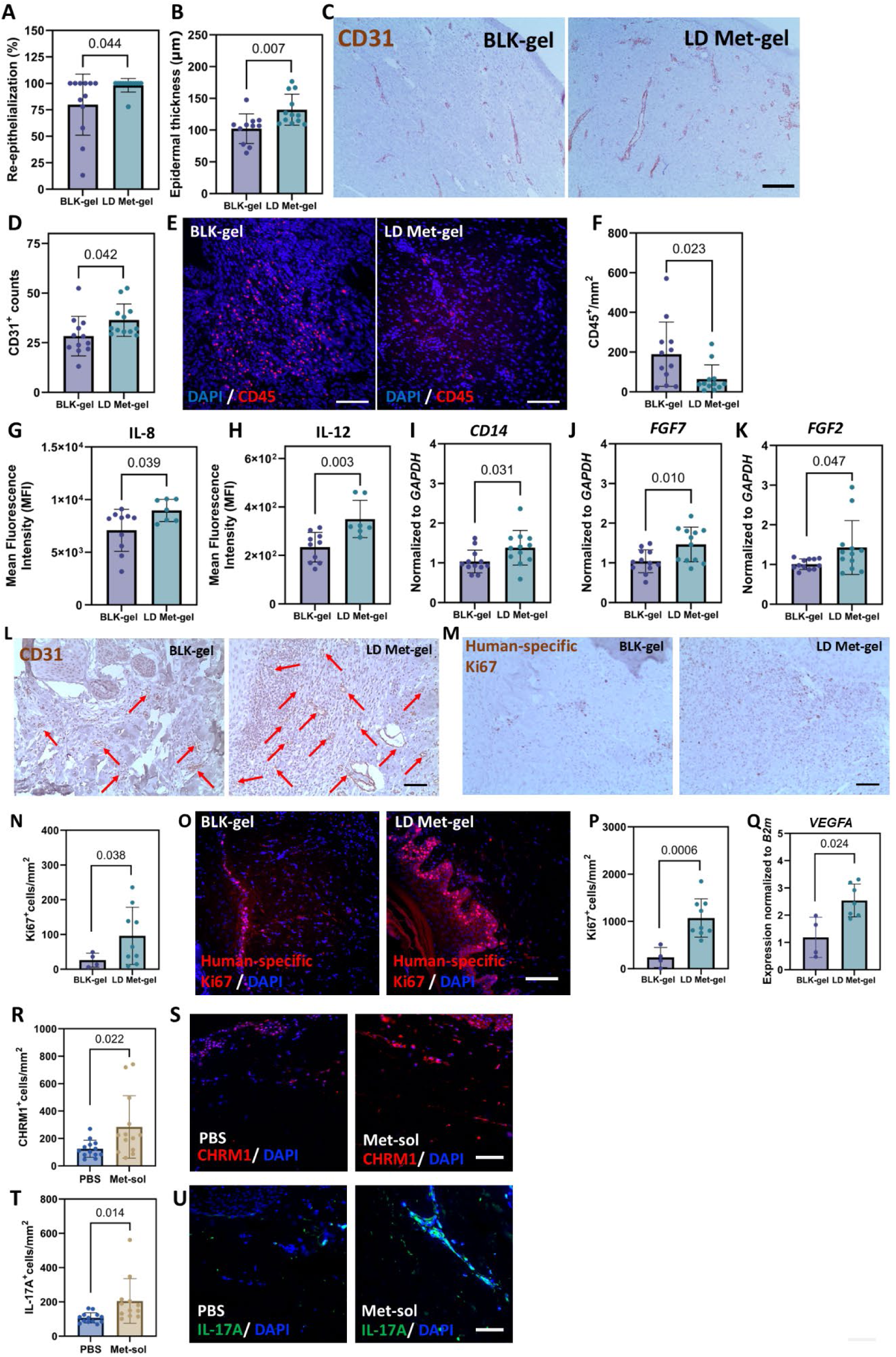
Metrifonate treatment promotes wound repair in diabetic porcine, human skin xenograft and ex vivo human skin models. **A,** Quantification of pig wounds re-epithelialization on Day (D) 21 expressed as %. **B**, Epidermal thickness of pig wounds on D21. **C**, Quantification of CD31+ vessels of pig wounds on D21. **D,** Representative immunohistochemistry (IHC) images on D21 for CD31, Scale bar=100 um **E,** Representative images of CD45 immunofluorescent staining in Met-gel and BLK gel treated diabetic porcine wounds. Scale bar=100um. **F,** Quantification of CD45+ cells per unit area. Data in **A-C** and **F** are presented as mean ± s.d. (*n* = 12). **G, H, Mean Fluorescence Intensity** (MFI) of IL-8 **(G)** and IL-12 **(H)** from porcine cytokine and chemokines Luminex assay. **I-K,** RT-qPCR gene expression analysis of porcine wounds on D21 for *CD14* **(I)**, *FGF7* **(J)**, and *FGF2* **(K)**. Data in **G-K** presented as mean ± s.d. (*n* = 12 for **I, J, K**; *n*=10 for BLK-gel, and *n*=7 for Met-gel for **G** and **H**). **L,** Representative IHC wound images on D7 of human skin engrafted diabetic mouse for CD31; Red arrows denote blood vessels, Scale bar=100 um. **M,** Representative wound IHC images on D7 for Ki67+ human cells in dermis, Scale bar=100 um. **N,** Quantification of Ki67+ human cells from dermis on D7 per unit area. **O,** Representative immunofluorescence wound images on D7 for Ki67+ human cells in epidermis, Scale bar=100 um. **P,** Quantification of Ki67+human cells from epidermis on D7 per unit area. **Q,** RT-qPCR gene expression analysis of wound on D7 for *VEGFA*. Data in **N, P, Q** presented as mean ± s.d. (*n* = 4 for BLK-gel, and *n*=7-10 for Met-gel). **R-U,** Quantification of CHRM1+ cells **(R)**, and IL-17A+ cells **(T)** from *ex vivo* human skin wounds on D4 per unit area. Data in **R, T** presented as mean ± s.d. (*n* = 12-14 wounds from 2 donors); Representative immunofluorescence image of CHRM1+cells **(S)** and IL-17A+ cells **(U)** from *ex vivo* human skin wound on D4. Scale bar= 100 um Statistical analyses were performed using two-sided unpaired *t*-tests as appropriate.

**Table S1.** Mouse Day 3 wound lysates multiplex cytokines results.

| **Analytes** | **LD Met-gel (pg/ml)** |  |  |  |  |  | **BLK-gel (pg/ml)** |  |  |  |  |  |
|---|---|---|---|---|---|---|---|---|---|---|---|---|
| **Eotaxin** | 135.74 | 251.16 | 183.64 | 226.35 | 198.59 | 319.2 | 164.49 | 208.14 | 231.29 | 241.92 | 189.89 | 178.61 |
| **6Ckine** | 942.67 | 1334.85 | 1119.28 | 895.45 | 1304.34 | 643.18 | 1549.04 | 821.53 | 761.94 | 786.33 | 858.8 | 920.79 |
| **Fractalkine** | 65.14 | 33.68 | 35 | 21.4 | 19.48 | 22.56 | 53.73 | 38.01 | 44.58 | 35.31 | 24.28 | 34.23 |
| **G-CSF** | 1352.78 | 5848.47 | 4592.4 | 4560.33 | 2139.62 | 6430.62 | 3071.62 | 4176.63 | 2251.97 | 1748.7 | 4500.63 | 5046.59 |
| **GM-CSF** | 7.91 | 11.94 | 10.47 | 10.84 | 13.56 | 11.7 | 5.59 | 5.25 | 8.2 | 6.09 | 13.9 | 8.34 |
| **IFNβ-1** | 59.27 | 120.15 | 10.55 | 29.25 | 40.17 | 19.91 | 15.5 | 22.46 | 84.04 | 92.73 | 61.71 | 29.91 |
| **IFNγ** | 8.78 | 2.44 | 3.55 | 3.6 | 3.9 | 3.4 | 2.18 | 1.16 | 3.95 | 1.98 | 3.3 | 2.69 |
| **IL-1α** | 924.82 | 555.52 | 491.64 | 671.6 | 684.7 | 558.05 | 624.76 | 320.85 | 579.92 | 838 | 268.79 | 646.48 |
| **IL-1β** | 7.74 | 15.94 | 28.59 | 33.54 | 9.51 | 21.58 | 15.37 | 15.15 | 14.92 | 20.19 | 26.24 | 18.12 |
| **IL-2** | 20.92 | 6.5 | 6.25 | 3.89 | 8.62 | 5.32 | 1.39 | 1.58 | 8 | 3.76 | 5.76 | 4.76 |
| **IL-3** | 0 | 0 | 0 | 0.19 | 0.11 | 0.51 | 0 | 0 | 0.14 | 0 | 0 | 0.09 |
| **IL-4** | 0.17 | 0.17 | 0.12 | 0.19 | 0.34 | 0.31 | 0.13 | 0.11 | 0.27 | 0.15 | 0.22 | 0.23 |
| **IL-5** | 0.11 | 0.13 | 0.25 | 0.45 | 0.16 | 0.37 | 0 | 0.11 | 0 | 0 | 0.18 | 0.11 |
| **IL-6** | 498.11 | 670.88 | 738.01 | 886.94 | 1709.4 | 6785.16 | 244.86 | 1061.47 | 219.12 | 928.24 | 430.56 | 1838.82 |
| **IL-7** | 0.85 | 0.46 | 0.75 | 0.09 | 0.65 | 0.85 | 0.27 | 0 | 0 | 0 | 0.09 | 0 |
| **IL-9** | 19.7 | 20.07 | 20.44 | 20.8 | 16.61 | 23.94 | 11.55 | 10.72 | 12.83 | 12.83 | 18.95 | 18.57 |
| **IL-10** | 10.64 | 15.93 | 3.93 | 8.97 | 8.14 | 4.49 | 38.59 | 19.83 | 7.58 | 2.79 | 19.83 | 7.02 |
| **IL-11** | 20.13 | 66.55 | 62.43 | 247.26 | 82.59 | 527.19 | 3.2 | 6.83 | 24.76 | 20.98 | 7.44 | 7.14 |
| **IL-12p70** | 6.23 | 9.87 | 5.62 | 14.46 | 9.26 | 14.15 | 20.58 | 78 | 40.08 | 111.61 | 491.88 | 254.37 |
| **IL-15** | 28.1 | 18.58 | 16.86 | 12.58 | 14.29 | 17.29 | 18.58 | 13.43 | 17.72 | 10.87 | 19.44 | 20.73 |
| **IL-16** | 154.36 | 254.65 | 183.44 | 142.81 | 142.03 | 150.11 | 86.15 | 2583.52 | 123.09 | 93.5 | 118.59 | 181.91 |
| **IL-17** | 4.5 | 2.61 | 4.55 | 13.49 | 11.36 | 17.09 | 1.09 | 1.26 | 3.56 | 1.56 | 5.83 | 5.89 |
| **IL-20** | 3.94 | 7.18 | 0 | 0.83 | 5.14 | 2.82 | 0 | 0.83 | 4.19 | 3.68 | 3.12 | 3.12 |
| **IP-10** | 8.82 | 12.1 | 18.28 | 14.81 | 20.92 | 47.11 | 9.26 | 11.76 | 11.28 | 19.86 | 21.57 | 15.89 |
| **KC** | 449.12 | 752.02 | 744.76 | 378.38 | 555.27 | 2501.3 | 330.72 | 419.26 | 757.42 | 618.05 | 1022.97 | 906.52 |
| **LIF** | 6.11 | 12.35 | 17.09 | 20.93 | 21.5 | 61.57 | 8.53 | 14.58 | 10.9 | 8.45 | 37.65 | 20.71 |
| **LIX** | 205.36 | 288.46 | 337.98 | 434.85 | 298.48 | 969.09 | 161.18 | 397.49 | 392.78 | 250.33 | 668.88 | 476.81 |
| **MCP-1** | 57.97 | 81.93 | 125.86 | 153.45 | 73.08 | 166.01 | 60.11 | 126.65 | 101.11 | 72.07 | 128.98 | 62.77 |
| **MCP-5** | 211.71 | 1153.33 | 325 | 470.93 | 193.89 | 226.52 | 514.4 | 987.57 | 220.27 | 220.09 | 269.14 | 363.41 |
| **MDC** | 14.94 | 19.14 | 12.6 | 16.89 | 24.88 | 25.93 | 6.89 | 28.39 | 10.98 | 17.92 | 19.8 | 16.12 |
| **M-CSF** | 10.99 | 9.2 | 19.42 | 12.13 | 13.42 | 8.12 | 9.98 | 9.91 | 20.42 | 14.99 | 12.06 | 7.83 |
| **MIG** | 19.55 | 20.91 | 91.65 | 65.79 | 111.81 | 59.16 | 15.16 | 23.32 | 11.58 | 18.53 | 57.81 | 39.71 |
| **MIP-1α** | 104.73 | 311.87 | 282.56 | 278.71 | 164.11 | 492.44 | 314.26 | 266.12 | 276.43 | 410 | 597.35 | 256.22 |
| **MIP-1β** | 58.71 | 336.74 | 281.44 | 331.98 | 152.95 | 561.1 | 175.36 | 178.87 | 165.59 | 238.7 | 516.87 | 309.85 |
| **MIP-2** | 4125.22 | 4614.53 | 10805.84 | 7578.9 | 4577.21 | 10281.31 | 6442.98 | 4293.02 | 7151.47 | 9706.1 | 5873.14 | 8445.87 |
| **MIP-3α** | 2.48 | 1.86 | 1.71 | 1.8 | 2.2 | 1.36 | 1.72 | 1.28 | 1.86 | 1.65 | 1.82 | 1.82 |
| **MIP-3β** | 43.04 | 40.86 | 31.08 | 47.71 | 37.58 | 26.55 | 92.04 | 78.13 | 29.35 | 70.36 | 18.62 | 38.98 |
| **RANTES** | 2.19 | 1.28 | 3.35 | 2.51 | 2.98 | 4.19 | 2.77 | 1.6 | 2.59 | 1.54 | 3.39 | 3.37 |
| **TARC** | 5.73 | 7.63 | 3.45 | 5.99 | 8.74 | 7.73 | 12.01 | 9.4 | 5.37 | 6.37 | 8.13 | 6.84 |
| **TNFα** | 3.39 | 19.38 | 32.88 | 34.38 | 14.41 | 26.46 | 79.74 | 29.39 | 13.41 | 22.16 | 29.04 | 20.2 |
| **VEGF** | 0.25 | 0.18 | 0.21 | 0.21 | 0.29 | 0.27 | 0.14 | 0.12 | 0.2 | 0.14 | 0.21 | 0.09 |

**Table S2.** Full list of genes and annotations for 10x Xenium panel.

| **Category – Cell Type** | **Gene** | **Comments** |
|---|---|---|
| NK cells / trNK | *Ncr1* | tr: tissue resident |
| NK cells / trNK | *Klrb1c* |  |
| NK cells / trNK | *Itga1* |  |
| NK cells / trNK | *Itga2* |  |
| NK cells / trNK | *Itgam* |  |
| NK cells / trNK | *Itgb2* |  |
| NK cells / trNK | *Gzma* |  |
| NK cells / trNK | *Prf1* |  |
| NK cells / trNK | *Il18r1* |  |
| NK cells / trNK | *Il18rap* |  |
| NK cells / trNK | *Tbx21* |  |
| NK cells / trNK | *Eomes* |  |
| NK cells / trNK | *Xcl1* |  |
| NK cells / trNK | *Klrg1* |  |
| NK cells / trNK | *Cd69* |  |
| NK cells / trNK | *Cxcr6* |  |
| NK cells / trNK | *Il7r* |  |
| NK cells / trNK | *Cd160* |  |
| NK cells / trNK | *Zeb2* |  |
| NK cells / trNK | *Il2rb* |  |
| NK cells / trNK | *Slamf7* |  |
| NK cells / trNK | *S1pr5* |  |
| NK cells / trNK | *Runx3* |  |
| NK cells / trNK | *Cd96* |  |
| NK cells / trNK | *Tnfsf10* |  |
| NK cells / trNK | *Emb* |  |
| NK cells / trNK | *Cxcr3* |  |
| NK cells / trNK | *Cxcr4* |  |
| NK cells / trNK | *Crtam* |  |
| NK cells / trNK | *Nr4a1* |  |
| NK cells / trNK | *Nr4a2* |  |
| NK cells / trNK | *Lmna* |  |
| NK cells / trNK | *Gpr183* |  |
| NK cells / trNK | *Ahr* |  |
| NK cells / trNK | *Ifng* |  |
| NK cells / trNK | *Klrc1* |  |
| NK cells / trNK | *Klrc2* |  |
| NK cells / trNK | *S100a4* |  |
| NK cells / trNK | *Cd226* |  |
| NK cells / trNK | *Sell* |  |
| NK cells / trNK | *Klra4* |  |
| NK cells / trNK | *Klra8* |  |
| NK cells / trNK | *Itga4* |  |
| NK cells / trNK | *Il15ra* |  |
| NK cells / trNK | *Kit* |  |
| NK cells / trNK | *Maf* |  |
| NK cells / trNK | *Inpp4b* |  |
| γδ T cells, iNKT, MAIT, Th17, Tc17 | *Cd3e* | IL-17 producing cells |
| γδ T cells, iNKT, MAIT, Th17, Tc17 | *Cd3g* |  |
| γδ T cells, iNKT, MAIT, Th17, Tc17 | *Cd4* |  |
| γδ T cells, iNKT, MAIT, Th17, Tc17 | *Cd8a* |  |
| γδ T cells, iNKT, MAIT, Th17, Tc17 | *Cd8b1* |  |
| γδ T cells, iNKT, MAIT, Th17, Tc17 | *Zbtb16* |  |
| γδ T cells, iNKT, MAIT, Th17, Tc17 | *Rorc* |  |
| γδ T cells, iNKT, MAIT, Th17, Tc17 | *Il17a* |  |
| γδ T cells, iNKT, MAIT, Th17, Tc17 | *Il17f* |  |
| γδ T cells, iNKT, MAIT, Th17, Tc17 | *Il23r* |  |
| γδ T cells, iNKT, MAIT, Th17, Tc17 | *Il1r1* |  |
| γδ T cells, iNKT, MAIT, Th17, Tc17 | *Mafa* |  |
| γδ T cells, iNKT, MAIT, Th17, Tc17 | *Mr1* |  |
| γδ T cells, iNKT, MAIT, Th17, Tc17 | *Cd1d1* |  |
| γδ T cells, iNKT, MAIT, Th17, Tc17 | *Il21* |  |
| γδ T cells, iNKT, MAIT, Th17, Tc17 | *Tcf7* |  |
| γδ T cells, iNKT, MAIT, Th17, Tc17 | *Pdcd1* |  |
| Cholinergic system | *Chrm1* | muscarinic/nicotinic receptors |
| Cholinergic system | *Chrm2* |  |
| Cholinergic system | *Chrm3* |  |
| Cholinergic system | *Chrm4* |  |
| Cholinergic system | *Chrm5* |  |
| Cholinergic system | *Chrna7* |  |
| Cholinergic system | *Chrna5* |  |
| Cholinergic system | *Chrnb2* |  |
| Cholinergic system | *Chrnb4* |  |
| Cholinergic system | *Ache* |  |
| Cholinergic system | *Chat* |  |
| Cholinergic system | *Slc5a7* |  |
| Cholinergic system | *Slc18a3* |  |
| Cholinergic system | *Rgs2* |  |
| Cholinergic system | *Grk2* |  |
| Cholinergic system | *Bche* |  |
| IL-17 signaling | *Il17ra* |  |
| IL-17 signaling | *Il17rc* |  |
| IL-17 signaling | *Ccl20* |  |
| IL-17 signaling | *Cxcl5* |  |
| IL-17 signaling | *Cxcl10* |  |
| IL-17 signaling | *Nfkb1* |  |
| IL-17 signaling | *Nfkb2* |  |
| IL-17 signaling | *Rela* |  |
| IL-17 signaling | *Il6* |  |
| IL-17 signaling | *Il1a* |  |
| IL-17 signaling | *Mmp3* |  |
| GM-CSF, IL-23, IL-6 axis | *Csf2* |  |
| GM-CSF, IL-23, IL-6 axis | *Csf2ra* |  |
| GM-CSF, IL-23, IL-6 axis | *Csf2rb* |  |
| GM-CSF, IL-23, IL-6 axis | *Il23a* |  |
| GM-CSF, IL-23, IL-6 axis | *Il12b* |  |
| GM-CSF, IL-23, IL-6 axis | *Il6ra* |  |
| GM-CSF, IL-23, IL-6 axis | *Il6st* |  |
| GM-CSF, IL-23, IL-6 axis | *Jak1* |  |
| GM-CSF, IL-23, IL-6 axis | *Stat3* |  |
| GM-CSF, IL-23, IL-6 axis | *Socs3* |  |
| T cells | *Cd3d* |  |
| T cells | *Ptprcap* |  |
| T cells | *Cd52* |  |
| T cells | *Ms4a4b* |  |
| T cells | *Ctsw* |  |
| T cells | *Rac2* |  |
| T cells | *Cd7* |  |
| T cells | *Hcst* |  |
| T cells | *Ltb* |  |
| T cells | *Cd2* |  |
| T cells | *Icos* |  |
| T regs | *Tnfrsf4* |  |
| T regs | *Foxp3* |  |
| T regs | *Il2ra* |  |
| T regs | *Ctla4* |  |
| Activated T cells | *Clu* |  |
| Cycling T cells | *Cks1b* |  |
| Dendritic cells | *Tap1* |  |
| Dendritic cells | *Itgax* |  |
| Dendritic cells | *Cd83* |  |
| Dendritic cells | *Cd86* |  |
| Dendritic cells | *Ccr7* |  |
| Dendritic cells | *Flt3* |  |
| Dendritic cells | *Tnfsf4* |  |
| Dendritic cells | *Irf8* |  |
| Dendritic cells | *Naga* |  |
| Dendritic cells | *Batf3* |  |
| Dendritic cells | *Ccl22* |  |
| Dendritic cells | *Cd209d* |  |
| Dendritic cells | *Cd209a* |  |
| Dendritic cells | *Fscn1* |  |
| Dendritic cells | *Ly6d* |  |
| Dendritic cells | *H2-DMa* |  |
| Dendritic cells | *Lsp1* |  |
| Dendritic cells | *Tbc1d4* |  |
| Dendritic cells | *Ifi30* |  |
| Dendritic cells | *Siglech* |  |
| Dendritic cells | *Cd207* |  |
| Dendritic cells | *Cxcl16* |  |
| Dendritic cells | *Nr4a3* |  |
| Dendritic cells | *Rel* |  |
| Dendritic cells | *Bcl2a1d* |  |
| Dendritic cells | *Cfp* |  |
| Dendritic cells | *Alox5ap* |  |
| Dendritic cells | *Pmaip1* |  |
| Monocytes / Macrophages | *Ly6c1* |  |
| Monocytes / Macrophages | *Ly6c2* |  |
| Monocytes / Macrophages | *Cytip* |  |
| Monocytes / Macrophages | *Fcgr1* |  |
| Monocytes / Macrophages | *Adgre1* |  |
| Monocytes / Macrophages | *Ccr2* |  |
| Monocytes / Macrophages | *Marco* |  |
| Monocytes / Macrophages | *Mertk* |  |
| Monocytes / Macrophages | *Cd68* |  |
| Monocytes / Macrophages | *Il10* |  |
| Monocytes / Macrophages | *Tgfbr1* |  |
| Monocytes / Macrophages | *Arg1* |  |
| Monocytes / Macrophages | *Nos2* |  |
| Monocytes / Macrophages | *Chil3* |  |
| Monocytes / Macrophages | *Ifitm6* |  |
| Monocytes / Macrophages | *Cx3cr1* |  |
| Monocytes / Macrophages | *Cd300c2* |  |
| Monocytes / Macrophages | *Mcub* |  |
| Monocytes / Macrophages | *Acp5* |  |
| Monocytes / Macrophages | *C1qb* |  |
| Monocytes / Macrophages | *Ctss* |  |
| Monocytes / Macrophages | *Csf1r* |  |
| Monocytes / Macrophages | *Selenop* |  |
| Monocytes / Macrophages | *Cbr2* |  |
| Monocytes / Macrophages | *C1qc* |  |
| Monocytes / Macrophages | *Ptgs2* |  |
| Monocytes / Macrophages | *Ccl8* |  |
| Monocytes / Macrophages | *Igf1* |  |
| Monocytes / Macrophages | *Gpnmb* |  |
| Monocytes / Macrophages | *Pf4* |  |
| Monocytes / Macrophages | *Lpl* |  |
| Monocytes / Macrophages | *Cd36* |  |
| Monocytes / Macrophages | *Fabp5* |  |
| Monocytes / Macrophages | *Pdpn* |  |
| Monocytes / Macrophages | *Plac8* |  |
| Monocytes / Macrophages | *Spic* |  |
| Monocytes / Macrophages | *Mrc1* |  |
| Monocytes / Macrophages | *Cd163* |  |
| Monocytes / Macrophages | *C1qa* |  |
| Monocytes / Macrophages | *Slc7a11* |  |
| Monocytes / Macrophages | *Cstdc4* |  |
| Monocytes / Macrophages | *Il1f9* |  |
| Monocytes / Macrophages | *Hcar2* |  |
| Monocytes / Macrophages | *Asprv1* |  |
| Monocytes / Macrophages | *Stfa2l1* |  |
| Monocytes / Macrophages | *Trem1* |  |
| Monocytes / Macrophages | *Arg2* |  |
| Monocytes / Macrophages | *C5ar1* |  |
| Monocytes / Macrophages | *Wfdc21* |  |
| Neutrophils | *Cxcr2* |  |
| Neutrophils | *Mmp9* |  |
| Neutrophils | *Elane* |  |
| Neutrophils | *Camp* |  |
| Neutrophils | *Lcn2* |  |
| Neutrophils | *Ngp* |  |
| Neutrophils | *Cebpe* |  |
| Neutrophils | *Csf3r* |  |
| Neutrophils | *Fpr1* |  |
| Neutrophils | *Lrg1* |  |
| Neutrophils | *Il1r2* |  |
| Neutrophils | *Egr1* |  |
| Neutrophils | *Egr2* |  |
| Neutrophils | *Rsad2* |  |
| Neutrophils | *Ifit1* |  |
| Neutrophils | *Ifit3* |  |
| Neutrophils | *Ly6g* |  |
| Neutrophils | *Isg20* |  |
| Neutrophils | *Slfn1* |  |
| Neutrophils | *Ccl9* |  |
| Neutrophils | *Lilrb4a* |  |
| Neutrophils | *Pglyrp1* |  |
| Basophils | *Mcpt8* |  |
| ILCs | *Tmem176a* | innate lymphoid cells |
| ILCs | *Ramp1* |  |
| ILCs | *Rora* |  |
| ILCs | *Il22* |  |
| ILCs | *Tox* |  |
| ILCs | *Nkg7* |  |
| ILCs | *Gata3* |  |
| ILCs | *Nfil3* |  |
| Mast Cells | *Ms4a2* |  |
| Mast Cells | *Cpa3* |  |
| Mast Cells | *Gata2* |  |
| Schwann Cells | *Matn2* |  |
| Schwann Cells | *Cavin2* |  |
| Schwann Cells | *Etv1* |  |
| Schwann Cells | *Des* |  |
| Schwann Cells | *Cd82* |  |
| Schwann Cells | *Ncmap* |  |
| Schwann Cells | *Pllp* |  |
| Melanocytes | *Dct* |  |
| Melanocytes | *Mlana* |  |
| Lymphatic Endothelial | *Lyve1* |  |
| Lymphatic Endothelial | *Cldn5* |  |
| PLC/Ca<sup>2+</sup> signaling | *Plcb1* | downstream of CHRM1 |
| PLC/Ca<sup>2+</sup> signaling | *Plcb3* |  |
| PLC/Ca<sup>2+</sup> signaling | *Plcb4* |  |
| PLC/Ca<sup>2+</sup> signaling | *Dagla* |  |
| PLC/Ca<sup>2+</sup> signaling | *Plcg1* |  |
| PLC/Ca<sup>2+</sup> signaling | *Plcg2* |  |
| PLC/Ca<sup>2+</sup> signaling | *Itpr1* |  |
| PLC/Ca<sup>2+</sup> signaling | *Plcd1* |  |
| PLC/Ca<sup>2+</sup> signaling | *Prkca* |  |
| PLC/Ca<sup>2+</sup> signaling | *Prkcb* |  |
| Calcium-regulated transcription | *Camk2a* | downstream of CHRM1 |
| Calcium-regulated transcription | *Camk2d* |  |
| Calcium-regulated transcription | *Nfatc1* |  |
| Calcium-regulated transcription | *Nfatc2* |  |
| Calcium-regulated transcription | *Nfatc3* |  |
| Calcium-regulated transcription | *Calm1* |  |
| Calcium-regulated transcription | *Stim1* |  |
| Calcium-regulated transcription | *Orai1* |  |
| PKC signaling effectors | *Mapk1* | downstream of CHRM1 |
| PKC signaling effectors | *Mapk3* |  |
| PKC signaling effectors | *Mapk14* |  |
| PKC signaling effectors | *Mapk8* |  |
| PKC signaling effectors | *Mapk9* |  |
| PI3K/Akt pathway | *Pik3ca* | downstream of CHRM1 |
| PI3K/Akt pathway | *Akt1* |  |
| PI3K/Akt pathway | *Rps6kb1* |  |
| VEGF and angiogenesis | *Vegfa* |  |
| VEGF and angiogenesis | *Vegfb* |  |
| VEGF and angiogenesis | *Vegfc* |  |
| VEGF and angiogenesis | *Flt1* |  |
| VEGF and angiogenesis | *Kdr* |  |
| VEGF and angiogenesis | *Nrp1* |  |
| VEGF and angiogenesis | *Angpt1* |  |
| VEGF and angiogenesis | *Angpt2* |  |
| VEGF and angiogenesis | *Tek* |  |
| Wound Healing Mediators | *Tgfb1* |  |
| Wound Healing Mediators | *Tgfb2* |  |
| Wound Healing Mediators | *Tgfb3* |  |
| Wound Healing Mediators | *Fn1* |  |
| Wound Healing Mediators | *Itgb1* |  |
| Wound Healing Mediators | *Vcam1* |  |
| Wound Healing Mediators | *Icam1* |  |
| Wound Healing Mediators | *Egf* |  |
| Wound Healing Mediators | *Fgf2* |  |
| Wound Healing Mediators | *Fgf7* |  |
| Wound Healing Mediators | *Hgf* |  |
| Wound Healing Mediators | *Pdgfa* |  |
| Wound Healing Mediators | *Pdgfb* |  |
| Wound Healing Mediators | *Ccn2* |  |
| Wound Healing Mediators | *Edn1* |  |
| Wound Healing Mediators | *Mmp2* |  |
| Wound Healing Mediators | *Timp1* |  |
| Wound Healing Mediators | *Timp2* |  |
| Wound Healing Mediators | *Nos3* |  |
| Wound Healing Mediators | *Pecam1* |  |
| Wound Healing Mediators | *Cav1* |  |
| Wound Healing Mediators | *Lox* |  |
| Wound Healing Mediators | *Areg* |  |
| Wound Healing Mediators | *Cxcl12* |  |
| Wound Healing Mediators | *Cd44* |  |
| Inflammation and Resolution | *Tnf* |  |
| Inflammation and Resolution | *Ccl5* |  |
| Inflammation and Resolution | *Cxcl9* |  |
| Inflammation and Resolution | *Nlrp3* |  |
| Inflammation and Resolution | *Pycard* |  |
| Inflammation and Resolution | *Casp1* |  |
| Inflammation and Resolution | *Socs1* |  |
| Inflammation and Resolution | *Irf1* |  |
| Inflammation and Resolution | *Irf5* |  |
| Inflammation and Resolution | *Irf7* |  |
| Inflammation and Resolution | *Stat1* |  |
| Tissue Remodeling and ECM | *Mmp8* |  |
| Tissue Remodeling and ECM | *Mmp13* |  |
| Tissue Remodeling and ECM | *Serpine1* |  |
| Tissue Remodeling and ECM | *Col4a1* |  |
| Tissue Remodeling and ECM | *Col5a1* |  |
| Tissue Remodeling and ECM | *Col6a1* |  |
| Tissue Remodeling and ECM | *Loxl1* |  |
| Tissue Remodeling and ECM | *Loxl2* |  |
| Tissue Remodeling and ECM | *Thbs1* |  |
| Tissue Remodeling and ECM | *Postn* |  |
| Tissue Remodeling and ECM | *Adam8* |  |
| Tissue Remodeling and ECM | *Adam10* |  |
| Tissue Remodeling and ECM | *Adamts1* |  |
| Tissue Remodeling and ECM | *Plaur* |  |
| Tissue Remodeling and ECM | *Plau* |  |
| Tissue Remodeling and ECM | *Serpinb2* |  |
| Tissue Remodeling and ECM | *Ptx3* |  |
| Stress/Metabolism/Immunometabolism | *Hif1a* |  |
| Stress/Metabolism/Immunometabolism | *Hmox1* |  |
| Stress/Metabolism/Immunometabolism | *Nfe2l2* |  |
| Stress/Metabolism/Immunometabolism | *Sod2* |  |
| Stress/Metabolism/Immunometabolism | *Gpx1* |  |
| Stress/Metabolism/Immunometabolism | *Gpx4* |  |
| Stress/Metabolism/Immunometabolism | *Cat* |  |
| Stress/Metabolism/Immunometabolism | *G6pdx* |  |
| Stress/Metabolism/Immunometabolism | *Slc2a1* |  |
| Stress/Metabolism/Immunometabolism | *Ldha* |  |
| Stress/Metabolism/Immunometabolism | *Pdk1* |  |
| Stress/Metabolism/Immunometabolism | *Glul* |  |
| Stress/Metabolism/Immunometabolism | *Acly* |  |
| Stress/Metabolism/Immunometabolism | *Mtor* |  |
| Stress/Metabolism/Immunometabolism | *Rptor* |  |
| Stress/Metabolism/Immunometabolism | *Prkaa1* |  |
| Stress/Metabolism/Immunometabolism | *Ddit3* |  |
| Proliferation | *Mki67* |  |
| Proliferation | *Pcna* |  |
| Proliferation | *Top2a* |  |
| Keratinocytes | *Aldh3a1* |  |
| Keratinocytes | *Sostdc1* |  |
| Keratinocytes | *Col17a1* |  |
| Keratinocytes | *Fcgbp* |  |
| Keratinocytes | *Krt75* |  |
| Keratinocytes | *Tm4sf1* |  |
| Keratinocytes | *Stmn1* |  |
| Keratinocytes | *Pclaf* |  |
| Keratinocytes | *Cdca8* |  |
| Keratinocytes | *Cdc20* |  |
| Keratinocytes | *Krt79* |  |
| Keratinocytes | *Defb6* |  |
| Keratinocytes | *Ldhb* |  |
| Keratinocytes | *Scd1* |  |
| Keratinocytes | *Krt1* |  |
| Keratinocytes | *Krt71* |  |
| Keratinocytes | *Krt25* |  |
| Keratinocytes | *Krt35* |  |
| Keratinocytes | *Crym* |  |
| Keratinocytes | *Fbp1* |  |
| Keratinocytes | *Kitl* |  |
| Keratinocytes | *Cxcl14* |  |
| Keratinocytes | *Ccl27a* |  |
| Keratinocytes | *Fgfbp1* |  |
| Keratinocytes | *Hspb1* |  |
| Keratinocytes | *Dnajb1* |  |
| Keratinocytes | *Perp* |  |
| Keratinocytes | *Krtdap* |  |
| Keratinocytes | *Cst6* |  |
| Keratinocytes | *Calm4* |  |
| Keratinocytes | *S100a14* |  |
| Keratinocytes | *Fosl1* |  |
| Basal Keratinocytes Proliferating | *Il4* |  |
| Basal Keratinocytes Proliferating | *Cyp11a1* |  |
| Basal Keratinocytes Proliferating | *Fcer1a* |  |
| Basal Keratinocytes Proliferating | *Cd200r3* |  |
| Basal Keratinocytes Proliferating | *Sytl3* |  |
| Basal Keratinocytes Proliferating | *Nusap1* |  |
| Basal Keratinocytes Proliferating | *Knl1* |  |
| Basal Keratinocytes Proliferating | *Aurka* |  |
| Basal Keratinocytes Proliferating | *Aurkb* |  |
| Basal Keratinocytes Proliferating | *Hmmr* |  |
| Basal Keratinocytes Proliferating | *Pbk* |  |
| Hair Follicle | *Cdh15* |  |
| Hair Follicle | *Aadac* |  |
| Hair Follicle | *Akr1c18* |  |
| Hair Follicle | *Car12* |  |
| Hair Follicle | *Prxl2a* |  |
| Pericytes | *Rps17* |  |
| Pericytes | *Selenow* |  |
| Pericytes | *Cavin3* |  |
| Pericytes | *Birc5* |  |
| Pericytes | *Ube2c* |  |
| Pericytes | *Pcp4l1* |  |
| Pericytes | *Lmod1* |  |
| Myofibroblast | *Tagln* |  |
| Myofibroblast | *Gm13889* |  |
| Myofibroblast | *Rgs16* |  |
| Myofibroblast | *Serpine2* |  |
| Myofibroblast | *Sparcl1* |  |
| Myofibroblast | *Tinagl1* |  |
| Endothelial Cells | *Rack1* |  |
| Endothelial Cells | *Meg3* |  |
| Endothelial Cells | *Aqp1* |  |
| Endothelial Cells | *Egfl7* |  |
| Endothelial Cells | *Ecscr* |  |
| Endothelial Cells | *Emcn* |  |
| Endothelial Cells | *Esam* |  |
| Endothelial Cells | *Selp* |  |
| Endothelial Cells | *Tmem252* |  |
| Endothelial Cells | *Cdh5* |  |
| Endothelial Cells | *Gpihbp1* |  |
| Endothelial Cells | *Col6a3* |  |
| SMC | *Mylk* | smooth muscle cell |
| SMC | *Actg2* |  |
| Fibroblasts | *Lum* |  |
| Fibroblasts | *Crabp1* |  |
| Fibroblasts | *Col7a1* |  |
| Fibroblasts | *Col14a1* |  |
| Fibroblasts | *Mgp* |  |
| Fibroblasts | *Pcolce2* |  |
| Fibroblasts | *Ndufa4l2* |  |
| Fibroblasts | *Col5a3* |  |
| Fibroblasts | *Prss23* |  |
| Fibroblasts | *Igfbp2* |  |
| Fibroblasts | *Megf6* |  |
| Fibroblasts | *Igfbp3* |  |
| Fibroblasts | *Bmp4* |  |
| Fibroblasts | *Coch* |  |
| Fibroblasts | *Dkk2* |  |
| Fibroblasts | *Dpp4* |  |
| Fibroblasts | *Lef1* |  |
| Fibroblasts | *Prdm1* |  |
| Fibroblasts | *Prss35* |  |
| Fibroblasts | *Runx1* |  |
| Fibroblasts | *Cnn1* |  |
| Fibroblasts | *Corin* |  |
| Fibroblasts | *Pappa2* |  |
| Fibroblasts | *Notum* |  |
| Fibroblasts | *Plin2* |  |
| Fibroblasts | *Wnt2* |  |
| Fibroblasts | *Fst* |  |
| Fibroblasts | *Twist2* |  |
| Fibroblasts | *Dlk1* |  |
| Fibroblasts | *Fmo1* |  |
| Fibroblasts | *Ly6a* |  |
| Fibroblasts | *Mest* |  |
| Fibroblasts | *Prg4* |  |
| Fibroblasts | *Col25a1* |  |
| Fibroblasts | *Pamr1* |  |
| Fibroblasts | *Fstl1* |  |
| Fibroblasts | *Bgn* |  |
| Fibroblasts | *Col16a1* |  |
| Fibroblasts | *Nupr1* |  |
| Fibroblasts | *Cd34* |  |
| Fibroblasts | *Ctsk* |  |
| Fibroblasts | *Aebp1* |  |
| Fibroblasts | *Eln* |  |
| Fibroblasts | *Mfap5* |  |
| Fibroblasts | *Cthrc1* |  |
| Fibroblasts | *Sfrp2* |  |
| Fibroblasts | *Tnfaip6* |  |
| Fibroblasts | *Dpt* |  |

**Table S3.** Porcine Day 4 wound exudates multiplex cytokines results.

| Analytes | LD Met-gel (pg/ml) |  |  |  |  |  |  | BLK-gel (pg/ml) |  |  |  |  |  |  |  |  |  |
|---|---|---|---|---|---|---|---|---|---|---|---|---|---|---|---|---|---|
| GM-CSF | OOR < | OOR < | 6.12 | OOR < | OOR < | 9.80 | OOR < | OOR < | OOR < | 119.77 | 55.47 | 85.14 | OOR < | OOR < | 1.39 | OOR < | OOR < |
| IFNγ | 96.93 | 122.07 | 273.91 | 159.79 | 147.21 | 248.34 | 147.21 | 71.76 | 96.93 | 1,226.15 | 432.41 | 736.81 | 96.93 | 122.07 | 159.79 | 59.16 | 96.93 |
| IL-1α | 3,936.91 | 3,056.00 | 3,738.93 | 5,520.74 | 4,340.31 | 6,325.41 | 2,293.32 | 5,216.91 | 621.98 | 2,709.21 | 4,074.47 | 4,174.26 | 3,866.47 | 2,279.55 | 2,005.49 | 3,479.62 | 1,966.36 |
| IL-1β | 10,794.77 | 36,626.51 | 3,214.69 | 9,356.90 | 2,093.41 | 24,246.10 | 43,797.57 | 6,340.65 | 11,830.50 | 27,211.09 | 10,548.46 | 16,829.94 | 16,102.35 | 21,642.86 | 36,283.64 | 22,335.71 | 31,846.82 |
| IL-1ra | 4,834.75 | 4,568.03 | 2,474.50 | 2,158.48 | 1,401.42 | 8,611.48 | 5,911.86 | 2,002.49 | 2,252.06 | 2,109.58 | 4,377.34 | 1,588.20 | 2,299.46 | 2,444.21 | 9,232.58 | 6,692.39 | 7,357.53 |
| IL-2 | OOR | OOR | 1.29 | OOR | OOR | 3.70 | OOR | OOR | OOR | 29.53 | 4.90 | 16.97 | OOR | OOR | OOR | OOR | OOR |
| IL-4 | OOR | OOR | OOR | OOR | OOR | OOR | OOR | OOR | OOR | 157.42 | 43.97 | 108.92 | OOR | OOR | OOR | OOR | OOR |
| IL-6 | 1,619.28 | 1,429.39 | 912.40 | 1,420.84 | 854.43 | 1,772.04 | 1,493.27 | 950.68 | 686.83 | 704.13 | 1,326.34 | 1,312.82 | 1,304.75 | 839.66 | 1,235.61 | 2,250.19 | 1,391.17 |
| IL-8 | OOR > | OOR > | 18,979.15 | 17,945.04 | 13,448.70 | OOR > | 211,826.58 | 36,652.52 | 3,296.71 | 1,877.34 | 25,259.00 | 7,494.37 | 23,460.24 | 4,697.82 | 21,468.75 | 28,430.13 | 23,725.42 |
| IL-10 | OOR | OOR | 33.71 | OOR | 1.20 | 70.62 | 15.11 | OOR | OOR | 317.07 | 141.01 | 215.03 | OOR | OOR | OOR | OOR | 24.41 |
| IL-12 | 236.77 | 352.81 | 218.23 | 233.40 | 196.41 | 354.54 | 229.60 | 229.60 | 108.31 | 107.49 | 178.42 | 199.76 | 127.72 | 87.73 | 171.33 | 189.29 | 220.76 |
| IL-18 | 6,928.42 | 10,698.98 | 4,184.63 | 8,894.52 | 4,715.66 | 11,847.61 | 7,650.80 | 8,338.53 | 3,797.94 | 6,456.14 | 9,740.44 | 5,655.93 | 5,096.57 | 2,940.66 | 12,048.70 | 11,020.00 | 8,711.66 |
| TNFα | OOR | 12.95 | 54.78 | 32.06 | 43.48 | 39.68 | 39.68 | 123.62 | 51.03 | 178.94 | 47.26 | 101.02 | OOR | 200.34 | 1.56 | OOR | 20.59 |
| Analytes | LD Met-gel (MFI) |  |  |  |  |  |  | BLK-gel (MFI) |  |  |  |  |  |  |  |  |  |
| GM-CSF | 11 | 12 | 18 | 12 | 15 | 19 | 14 | 10 | 12 | 75 | 38 | 54 | 14 | 13 | 17 | 14 | 12 |
| IFNγ | 16 | 17 | 23 | 19 | 18 | 22 | 18 | 15 | 16 | 52 | 29 | 39 | 16 | 17 | 19 | 15 | 16 |
| IL-1α | 9,023 | 8,207 | 8,862 | 9,997 | 9,319 | 10,345 | 7,224 | 9,845 | 3,113 | 7,800 | 9,129 | 9,202 | 8,967 | 7,203 | 6,755 | 8,633 | 6,686 |
| IL-1β | 5,083 | 9,768 | 2,044 | 4,623 | 1,410 | 8,088 | 10,498 | 3,506 | 5,391 | 8,554 | 5,007 | 6,661 | 6,495 | 7,635 | 9,730 | 7,760 | 9,196 |
| IL-1ra | 1,824 | 1,760 | 1,152 | 1,039 | 741 | 2,623 | 2,067 | 981 | 1,073 | 1,021 | 1,712 | 819 | 1,090 | 1,141 | 2,744 | 2,233 | 2,371 |
| IL-2 | 13 | 14 | 20 | 18 | 16 | 22 | 13 | 17 | 13 | 43 | 23 | 33 | 16 | 14 | 14 | 14 | 15 |
| IL-4 | 16 | 19 | 21 | 19 | 20 | 22 | 20 | 19 | 15 | 46 | 30 | 39 | 19 | 19 | 17 | 18 | 18 |
| IL-6 | 1,875 | 1,748 | 1,331 | 1,742 | 1,275 | 1,970 | 1,792 | 1,366 | 1,101 | 1,120 | 1,674 | 1,664 | 1,658 | 1,261 | 1,606 | 2,230 | 1,721 |
| IL-8 | 10,068 | 9,928 | 8,070 | 8,011 | 7,659 | 10,062 | 9,030 | 8,578 | 4,667 | 3,164 | 9,106 | 6,665 | 8,269 | 5,595 | 8,190 | 8,418 | 8,278 |
| IL-10 | 19 | 17 | 23 | 17 | 20 | 27 | 21 | 15 | 13 | 57 | 35 | 44 | 19 | 16 | 19 | 19 | 22 |
| IL-12 | 323 | 459 | 301 | 319 | 275 | 461 | 315 | 315 | 169 | 168 | 254 | 279 | 193 | 144 | 245 | 267 | 304 |
| IL-18 | 710 | 999 | 460 | 868 | 511 | 1,076 | 770 | 825 | 421 | 670 | 931 | 599 | 547 | 333 | 1,089 | 1,021 | 854 |
| TNFα | 20 | 25 | 36 | 30 | 33 | 32 | 32 | 56 | 35 | 75 | 34 | 49 | 21 | 83 | 22 | 21 | 27 |

**Table S4.** List of primary antibodies for immunofluorescence and immunohistochemistry staining, and western blot analyses.

| Species Applied | Immunofluorescence |  |  |
|---|---|---|---|
| **Mouse** | **Antibody** | **Dilution** | **Product #** |
|  | rabbit anti-CD31 | 1:100 | ab28364 |
|  | rat anti-Ki67 mouse specific | 1:100 | HS-398 117 |
|  | rat anti-ROR gamma t | 1:100 | AFKJS-9 |
|  | rabbit anti-IL17A | 1:100 | bs-1183R |
|  | rabbit anti-Interleukin-18 Receptor Beta | 1:100 | CAU25549 |
|  | goat anti-NKp46/NCR1 | 1:100 | AF2225 |
|  | armenian hamster anti-TCR beta | 1:100 | H57-597 |
|  | armenian hamster anti-TCR γ/δ | 1:100 | 118101 |
|  | goat anti-Mcpt4 | 1:100 | aa114-126 |
|  | armenian hamster Anti-CD11c | 1:100 | ab33483 |
|  | rat Neutrophil Marker (NIMP-R14) | 1:100 | sc-59338 |
|  | rabbit anti-F4/80 [SP115] | 1:100 | ab111101 |
|  | rabbit anti-Klrb1c | 1:100 | bs-4682R |
|  | rabbit anti-CHRM1 | 1:100 | NBP1-87466 |
|  | mouse anti-Acetylcholinesterase [HR2] | 1:100 | ab2803 |
| **Human** | mouse anti-IL-17A | 1:100 | eBio64DEC17 |
|  | goat anti-NKp46/NCR1 | 1:100 | AF2225 |
|  | rabbit anti-Ki67 human specific | 1:100 | HS-398 003 |
|  | goat anti-CHRM1 | 1:100 | ab77098 |
|  | rabbit anti-CHRM1 | 1:100 | A04081 |
| **Porcine** | mouse anti-CD45 | 1:100 | K252.1E4 |
| **Species reactivity** | **Immunohistochemistry** |  |  |
| **Mouse** | **Antibody** | **Dilution** | **Product #** |
|  | rabbit polyconal anti-CD31 | 1:100 | ab28364 |
| **Human** | rabbit anti0 Ki67 human specific | 1:100 | HS-398 003 |
|  | rabbit anti-CD31 | 1:100 | ab28364 |
| **Porcine** | rabbit anti-CD31 | 1:100 | ab28364 |
|  | **Western Blot** |  |  |
|  | **Antibody** | **Dilution** | **Product #** |
| **Mouse** | rabbit anti-Lipocalin-2 / NGAL | 1:1000 | ab63929 |
|  | mouse anti-MMP9 | 1:1000 | sc-393859 |
|  | mouse anti-TIMP-1(2A5) | 1:1000 | sc-21734 |
|  | mouse anti-Vinculin (7F9) | 1:1000 | sc-73614 |

**Table S5.**
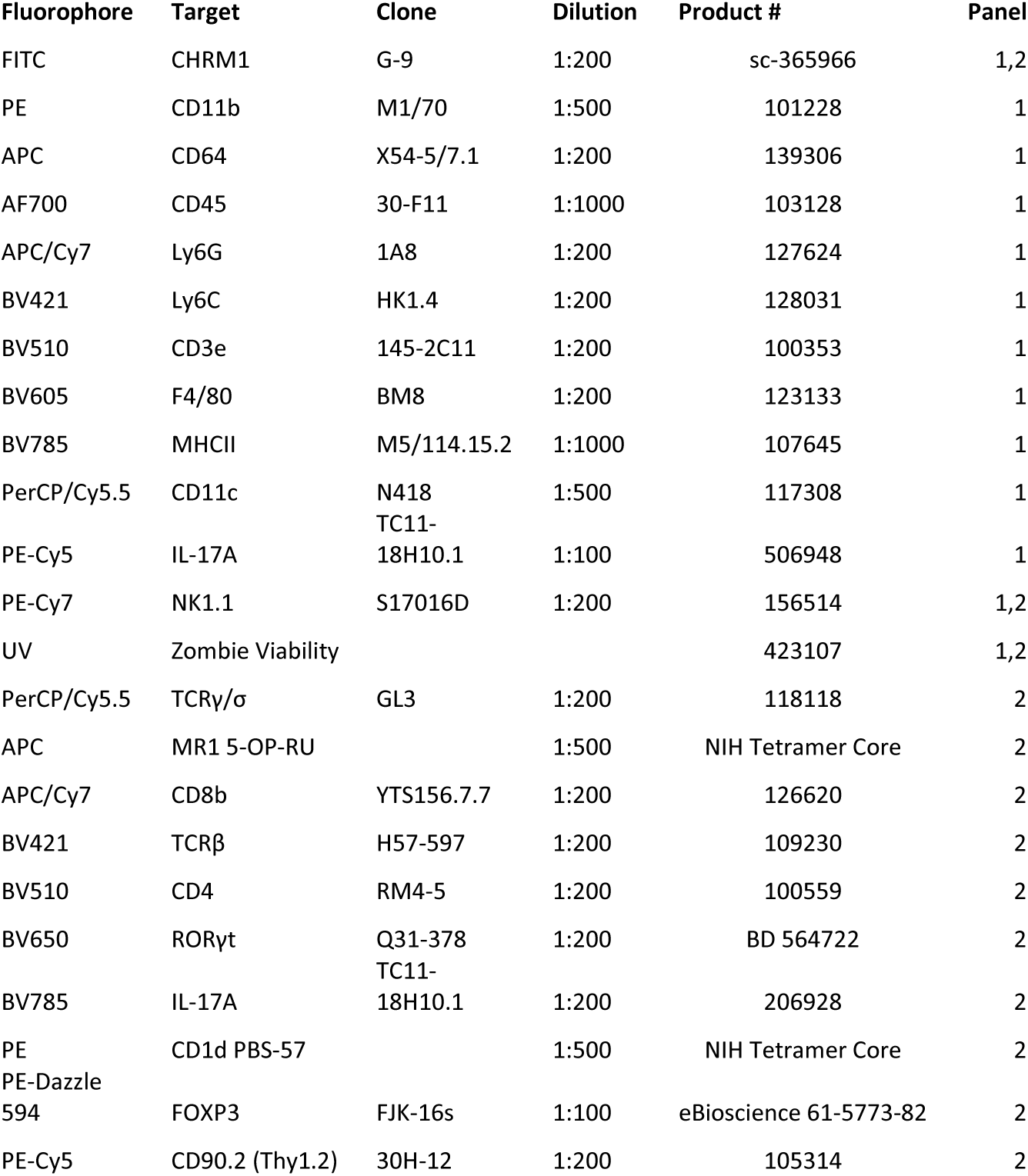
List of antibodies for flow cytometry panels. Antibodies were purchased from Biolegend unless otherwise listed.

**Table S6.** List of primers for RT-qPCR, purchased from QIAGEN.

| **Mouse Gene** | **GeneGlobe ID** |
|---|---|
| *B2m* | QT01149547 |
| *Actb* | QT00095242 |
| *Tgfb1* | QT00145250 |
| *Acta2* | QT00140119 |
| *Arg1* | QT00134288 |
| *Il10* | QT00106169 |
| *Chil3* | QT00108829 |
| *Il1b* | QT01048355 |
| *Mmp9* | QT00108815 |
| *Adipoq* | QT01048047 |
| *Fabp4* | QT00091532 |
| *B2m* | PPM03562A |
| *Actb* | PPM02945B |
| *Chrm1* | PPM03990A |
| *Chrm2* | PPM34851A |
| *Chrm3* | PPM03891B |
| *Chrm4* | PPM03988A |
| *Chrm5* | PPM38763A |
| *Chrna3* | PPM04298A |
| *Chrna4* | PPM03963A |
| *Chrna5* | PPM30677A |
| *Chrna7* | PPM03977A |
| *Chrnb2* | PPM04007A |
| *Chrnb4* | PPM27992A |
| **Human Gene** | **GeneGlobe ID** |
| *GAPDH* | QT00079247 |
| *VEGFA* | QT01010184 |
| **Pig Gene** | **GeneGlobe ID** |
| *CD14* | PPS01617A |
| *FGF2* | PPS00943A |
| *FGF7* | PPS00608A |

**Table S7.** Skin donor information.

| **Split-thickness human skin from BioIVT for xenotransplantation** |  |  |  |  |  |
| **Sex** | **Age** | **Anatomical site** | **Safety data** |  |  |
|---|---|---|---|---|---|
| F | 33 | Abdominoplasty | Negative for HepC, Hep B, HIV-1, and HIV-2 |  |  |
| **Ex vivo human skin from Genoskin** |  |  |  |  |  |
| **Donor ID** | **Age** | **Sex** | **BMI** | **Anatomical site** | **Safety data** |
| **1** | 48 | F | 30.2 | Abdominoplasty | Negative for Hep C, Hep B, HIV-1, and HIV-2 |
| **2** | 36 | F | 30.1 | Abdominoplasty | Negative for Hep C, Hep B, HIV-1, and HIV-2 |
| **3** | 35 | F | 26.8 | Abdominoplasty | Negative for Hep C, Hep B, HIV-1, and HIV-2 |

